# TBK1 activity regulates the directionality of axonal transport of signalling endosomes

**DOI:** 10.1101/2025.09.03.674015

**Authors:** David Villarroel-Campos, Jose Norberto S Vargas, Riccardo Zenezini Chiozzi, Anna-Leigh Brown, Martin Wallace, Kai Sun, James N. Sleigh, Konstantinos Thalassinos, Pietro Fratta, Giampietro Schiavo

## Abstract

The polarised and complex morphology of neurons pose massive challenges for efficient cargo delivery between the axon and soma, a process termed axonal transport. We have previously shown that the retrograde axonal transport of pro-survival, neurotrophic signalling endosomes relies on Rab7 in motor neurons, and that their trafficking is impaired in the early stages of amyotrophic lateral sclerosis (ALS) pathogenesis. Here, we report the effect of Rab7 phosphorylation on the transport of these signalling endosomes. We show that the ALS-linked kinase TBK1 phosphorylates Rab7 at S72 in neurons, altering its binding to cytoplasmic dynein adaptors. Accordingly, both TBK1 knockdown and the expression of a loss-of-function Rab7 mutant (S72E) induce aberrant bidirectional movement of signalling endosomes without modifying neuronal polarity or endosomal sorting. This alteration is specific for signalling endosomes, as axonal transport of lysosomes and mitochondria remain unaffected. We have therefore discovered a new TBK1 function that ensures the unidirectional transport of signalling endosomes, suggesting that reduced TBK1 activity determines retrograde transport dysfunctions and long-range signalling impairments.

**Summary:** Signalling endosomes travel from the distal axon to the cell body, exerting pro-survival functions. TBK1, an ALS-linked kinase, governs the directionality of signalling endosome transport in motor neurons by phosphorylating Rab7. Thus, loss of TBK1 function results in long-range signalling impairment.

## Introduction

The complex morphology of neurons, and their polarisation into somatodendritic and axonal compartments are essential for their function. The large distance encompassing these compartments, which in peripheral neurons can be more than a metre, raises formidable challenges to long-range intracellular communication. For instance, neuronal homeostasis relies on the bidirectional delivery of a variety of cargoes along the axon, a process termed axonal transport, which becomes deregulated in neurological disorders [1]. In particular, target tissues provide neurotrophic factors that are retrogradely transported from axon terminals, and exert pro-survival signalling in the cell body [2]. This transport occurs in specialised endocytic organelles known as signalling endosomes, which retain signalling competence during their journey back to the soma [3]. We have shown that brain-derived neurotrophic factor (BDNF) and its receptor tropomyosin receptor kinase B (TrkB) are retrogradely transported in signalling endosomes carrying Rab7A (hereafter referred as Rab7) [4], and that this process is impaired both *in vitro* and *in vivo* in genetic models of amyotrophic lateral sclerosis (ALS) [5–7].

Rab GTPases are crucial regulators of intracellular trafficking, coordinating the fission, transport, tethering and fusion of several cytoplasmic compartments [8]. They act as molecular switches, cycling through a GTP-bound active state and a GDP-bound inactive state, promoted by guanine nucleotide exchange factors (GEFs) and GTPase-activating proteins (GAPs), respectively. In addition, Rab function is regulated by post-translation modifications, such as phosphorylation [9]. In particular, Rab7 is phosphorylated at S72 (p-S72) by TANK-binding kinase 1 (TBK1), Leucine Rich Repeat Kinase 1 (LRRK1), Transforming growth factor beta-activated kinase 1 (TAK1), NIMA-related kinase 7 (NEK7), and Inhibitor of Nuclear Factor Kappa B kinase subunit epsilon (IKKε) [10–14]. S72 phosphorylation impairs the interaction between Rab7 and GDP dissociation inhibitor (GDI), and alters the recruitment of effectors [15], with some of them preferentially binding either phosphorylated or unphosphorylated Rab7 [16], thus impacting on Rab7 cellular functions.

TBK1 is a Ser/Thr kinase involved in the regulation of innate immunity and selective autophagy [17,18]. Heterozygous *TBK1* mutations lead to haploinsufficiency and cause ALS [19,20], although the mechanisms linking these mutations to ALS phenotypes remain disputed. Studies carried out in non-neuronal cells have shown that TBK1-dependent Rab7 p-S72 is required for PARKIN-dependent selective mitophagy [10], for the regulation of the immune response downstream to stimulator of interferon genes (STING) in breast cancer cells [14] and for relieving mammalian target of rapamycin complex 1 (mTORC1) inhibition in response to amino acid re-feeding in starved cells [21]. Rab7 GTPase activity is required for axonal transport of signalling endosomes, since expression of a Rab7 dominant-negative mutant severely affects their transport [4]. Given that the role of Rab7 phosphorylation in the trafficking of signalling endosomes and other cytoplasmic organelles in neurons remains unaddressed, we decided to test whether TBK1 regulates the retrograde transport of signalling endosomes in mouse motor neurons (MNs) through Rab7 phosphorylation.

Here, we report a new role for TBK1 in maintaining the unidirectional retrograde movement of signalling endosomes in primary MNs, through the regulation of dynein adaptors recruited by Rab7 p-S72. Since neurotrophic signals use the endocytic pathway to reach the soma, and the same pathway is also impinged by ALS-linked TBK1 deficiency [22], we demonstrate a novel function for TBK1 that may contribute to motor neuron pathology.

## Results

### TBK1 loss-of-function impairs the axonal transport of signalling endosomes

ALS-linked *TBK1* mutations are either nonsense or missense mutations, pointing to a loss-of-function pathomechanism [23,24]. We therefore decided to pursue a knockdown strategy, testing two shRNAs (shTBK1-1 and shTBK1-2) together with a scramble control (shControl) in wild-type (WT) mouse primary motor neuron cultures (**Supplementary Figure 1**). MNs transduced with shTBK1-1 or shTBK1-2 showed ∼ 65% knockdown after 72 h, whilst MNs transduced with shControl displayed TBK1 expression levels equivalent to untransduced neurons (**Supplementary Figure 1A,B**). TBK1 knockdown is greater when assessed by western blotting (**Supplementary Figure 1C,D**), likely due to the contribution of glial cells in the primary motor neuron culture. We also confirmed that TBK1 knockdown does not affect neuron viability at 6, 9 and 12 days *in vitro* (DIV) (**Supplementary Figure 1E,F**), in agreement with previous reports [25].

Having validated these two TBK1 shRNAs, we transduced primary MNs on DIV 3 and imaged axonal signalling endosomes three days later. For imaging purposes, these organelles were labelled with a non-toxic fragment of tetanus neurotoxin (H_C_T), which has been shown to undergo axonal retrograde transport in endocytic carriers shared with neurotrophin receptors [4,26]. Representative kymographs and displacement graphs for these conditions are shown in **Figure 1A** and **1B**, respectively. Most of the cargoes in untreated and shControl conditions move retrogradely. In contrast, a significant proportion of signalling endosomes exhibit anterograde transport upon TBK1 knockdown with each shRNA vector (**Figure 1B**). Indeed, the speed distribution profiles in both the shTBK1-1 and shTBK1-2 conditions are similarly shifted to the left (**Figure 1C**), reflecting an increase in anterograde transport. Interestingly, the net average speed per cargo remains unaffected (**Figure 1D**), therefore, this phenotype arises from a shift in directionality rather than from a change in the overall cargo speed. As expected, TBK1 knockdown using shTBK1-1 (hereafter called shTBK1) induces an accumulation of signalling endosomes at neurite tips (**Figure 1E**). Thus, loss of TBK1 in MNs leads to a population of signalling endosomes moving anterogradely, which fail to deliver their pro-survival signals to the soma.

**Figure 1.**
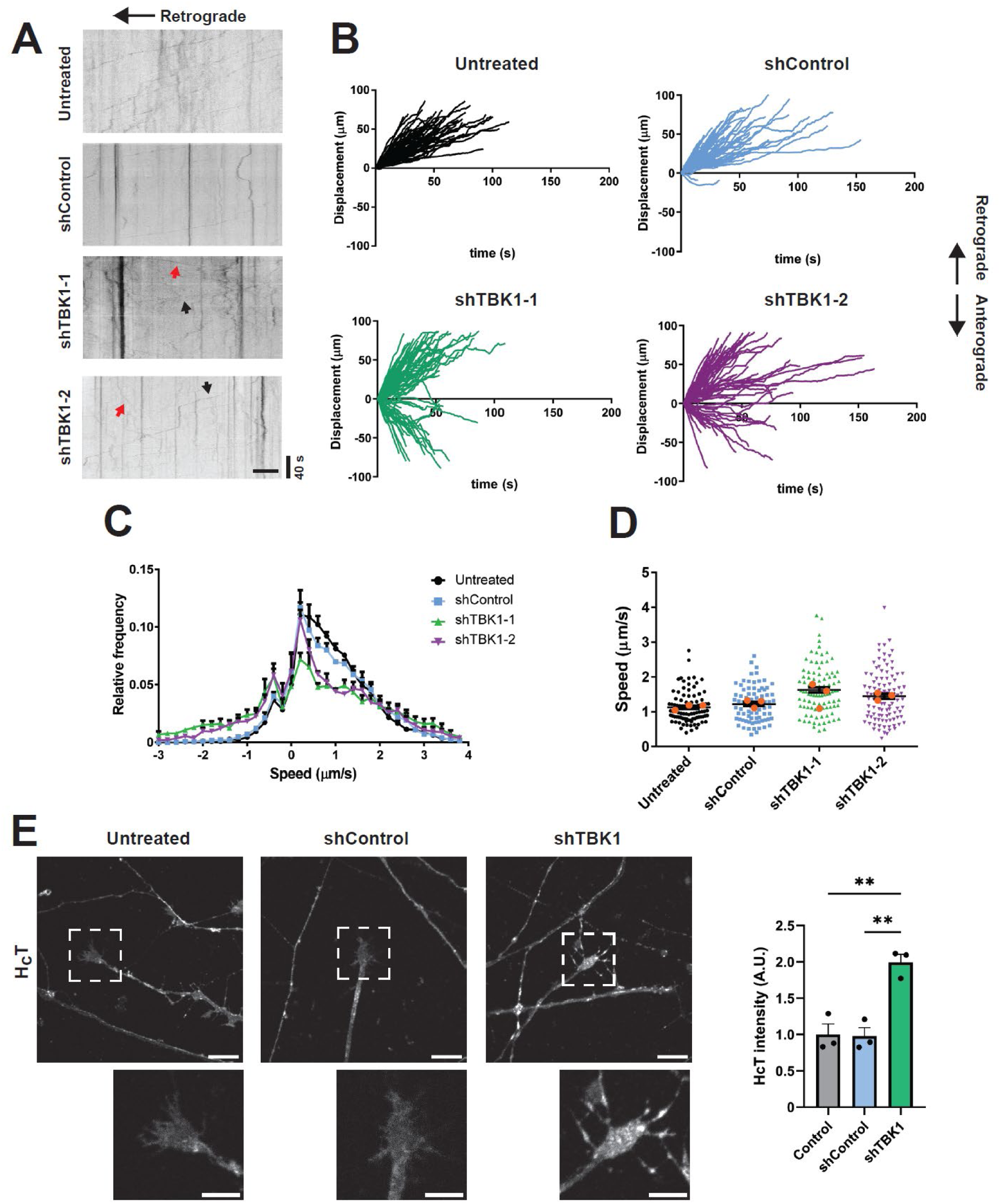
TBK1 knockdown induces bidirectional movement in a population of signalling endosomes. Primary MNs were transduced with lentiviral vectors encoding shControl, shTBK1-1 or shTBK1-2 on DIV 3. After 72 h, cells were labelled with 30 nM AlexaFluor 647-H_C_T for 45 min, washed and imaged. **(A)** Representative kymographs for all conditions, with the cell body located towards the left. In the shTBK1 conditions, black arrows indicate retrograde movement, whilst red arrows show anterograde movement. Scale bar, 10 μm. **(B)** Displacement graphs for all conditions, with retrograde transport shown as positive. A sub-population of carriers moves anterogradely for both shTBK1-1 and shTBK1-2. **(C)** Speed profiles for each condition; both shTBK1-1 and shTBK1-2 curves are shifted to the left, indicating increased anterograde movement. **(D)** Average speed per cargo, considering all displacement as positive. The average speed is not affected. Number of cargoes: Untreated (Unt) 92, shControl 79, shTBK1-1 92, shTBK1-2 103, from three independent cultures, orange dots represent the mean for each replicate; one-way ANOVA with Tukey’s multiple comparison test. **(E)** Primary MNs were labelled with 30 nM AlexaFluor 555-H_C_T for 45 min, washed and fixed. Images from neurite tips show an accumulation of H_C_T at distal regions in the shTBK1 condition. Scale bar, 10 μm in main panels and 5 μm in the insets. n = 3 different cultures; one-way ANOVA with Tukey’s multiple comparison test, **, P < 0.01.

### Rab7 S72E expression causes similar alterations in signalling endosome transport

Since TBK1 loss-of-function modifies the axonal movement of signalling endosomes, we next tested how mutations in Rab7 S72, a residue phosphorylated by TBK1, affect this process. We transfected GFP-Rab7 WT, S72A or S72E into DIV5 primary MNs using magnetofection, and imaged signalling endosomes labelled with H_C_T on DIV 6. In transfected neurons, we detected two populations of cargoes: 1) endosomes double positive for Rab7 and H_C_T, and 2) single positive endosomes labelled with H_C_T alone, except in the Rab7 S72E condition, where just H_C_T single positive cargoes were found, in agreement with the cytoplasmic localisation of the S72E mutant (**Supplementary Figure 2A**) [15]. Representative kymographs (**Figure 2A**) and displacement graphs (**Figure 2B**) show that most cargoes moved towards the soma, except for a subpopulation of signalling endosomes in the Rab7 S72E condition, which were transported anterogradely. In accordance with the displacement analysis, the speed distribution profiles for double positive or H_C_T-only cargoes in MNs expressing Rab7 S72A are indistinguishable from those expressing Rab7 WT (**Figure 2C,D**). However, single-positive signalling endosomes in MNs expressing Rab S72E display a significant shift to the left (**Figure 2D**), in agreement with the increase in anterograde transport shown in **Figure 2B**. The average speed per cargo, considering every displacement as positive, shows that cargoes move at similar speeds in every condition (**Figure 2E**). Altogether, these results indicate that Rab7 S72E disrupts the axonal transport of signalling endosomes by enabling a directionality switch similar to that observed upon TBK1 knockdown.

**Figure 2.**
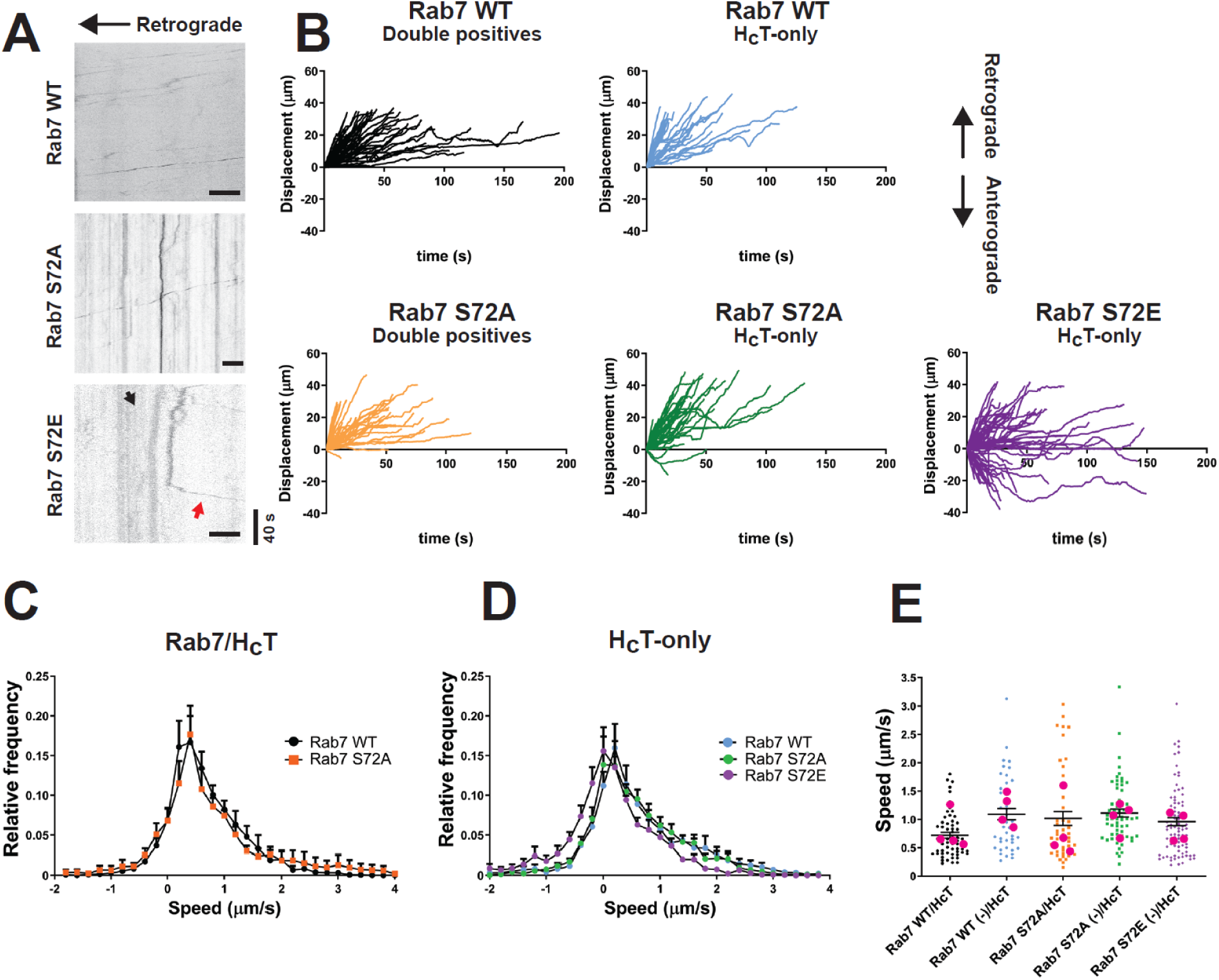
Rab7 S72E disrupts signalling endosome axonal transport. Primary MNs were magnetofected with GFP-Rab7 WT, S72A or S72E on DIV 5. 24 h later, signalling endosomes were labelled with 30 nM AlexaFluor 555-H_C_T for 45 min, washed and imaged. Cargoes were tracked and classified as double positive (GFP-Rab7 and H_C_T) or single positive (H_C_T-only). **(A)** Representative kymographs for all conditions, with the cell body located towards the left. In the S72E condition, the black arrow indicates retrograde movement, whilst the red arrow shows anterograde movement. Scale bar, 10 μm. **(B)** Displacement graphs for every condition, with retrograde movement considered as positive. H_C_T-containing cargoes move towards the soma, however a subpopulation of endosomes in the S72E condition moves anterogradely. **(C)** Speed profiles for Rab7/H_C_T cargoes. Their speed distribution is not changed. **(D)** Speed profiles for single positive cargoes. The Rab7 S72E curve is shifted to the left, reflecting an increase in anterograde movement. **(E)** Average speed per cargo, considering every displacement as positive. The speed of H_C_T-positive cargoes is not affected. Number of cargoes: WT/H_C_T 59, WT (-)/H_C_T 43, S72A/H_C_T 45, S72A (-)/H_C_T 62, S72E (-)/H_C_T 85 from four independent primary cultures, magenta dots represent the mean for each replicate; one-way ANOVA with Tukey’s multiple comparison test.

### The bidirectional transport of signalling endosomes is not linked to mis-sorting or polarity defects

Since both Rab7 S72E expression and TBK1 knockdown affect the directionality of transport of H_C_T-positive signalling endosomes, we asked whether this effect is due to alterations of neuronal polarity. We magnetofected primary MNs with Rab7 WT or S72E on DIV 2 and subsequently stained them for the somatodendritic marker MAP2 and the axonal marker SMI-31. As shown in **Supplementary Figure 2A**, our analysis did not reveal any overt abnormality in neuronal polarity in these neurons. In addition, we magnetofected primary MNs with these Rab7 variants and EB3-mCherry to visualise the polarity of axonal microtubules (**Supplementary Figure 2B**); we found that ∼ 98% of microtubules are oriented with their plus-ends facing the axonal tip in both conditions. To test whether Rab7 S72E impacts the axonal pattern of acetylated (Ac-Tub) and tyrosinated (Tyr-Tub) tubulin, we simultaneously extracted the soluble fraction of the cytoskeleton and fixed MNs expressing Rab7 WT, S72A or S72E (**Supplementary Figures 2C**). MNs expressing the different Rab7 variants exhibit a similar enrichment of Tyr-Tub at the axonal tip, and undistinguishable ratios of Tyr-Tub/Ac-Tub in the proximal and distal axon (**Supplementary Figure 2D**). Altogether, these experiments suggest that the observed axonal transport phenotype is not related to changes in neuronal polarity or structural alterations of the microtubule cytoskeleton.

Previous reports suggest that autophagosomes sequentially engage different dynein adaptors during their axonal transport towards the neuronal soma; as a consequence, their otherwise unidirectional movement becomes bidirectional upon maturation through fusion with lysosomes [27,28]. Therefore, we asked whether the observed directionality phenotype may arise from defective sorting of signalling endosomes into lysosomes. We transduced primary MNs on DIV 3 with shControl or shTBK1 and labelled signalling endosomes with H_C_T and lysosomes with lysotracker on DIV 6, to quantify double-positive organelles. We found organelles positive for both markers (**Supplementary Figure 2E**, arrows) and compartments labelled only with H_C_T (**Supplementary Figure 2E**, arrowheads). Further quantification showed that ∼ 25% of signalling endosomes were also labelled by lysotracker, a percentage that did not vary among treatments (**Supplementary Figure 2F**), ruling out mis-sorting to lysosomes as the main cause of bidirectional trafficking.

### TBK1- and Rab7 S72E-dependent bidirectional transport is signalling endosome-specific

Neurotrophin signalling endosomes share common features with late endosomes and lysosomes in spinal MNs, such as the presence of Rab7 in their limiting membrane [4]. Therefore, we wanted to determine whether the transport disruption caused by TBK1 downregulation also affects these related cargoes. We thus transduced primary MNs with shControl or shTBK1 on DIV 3, and tracked lysosomes labelled with lysotracker on DIV 6. In agreement with previous reports, we found that lysosomes move bidirectionally [29], with ∼ 80% of lysosomes moving towards the soma, a percentage that was unchanged across conditions (**Figure 3A,B**). Additionally, TBK1 knockdown has no effect on the speed profile of these organelles (**Figure 3C**) nor on their average speed (**Figure 3D**), compared to the untreated condition (Unt).

**Figure 3.**
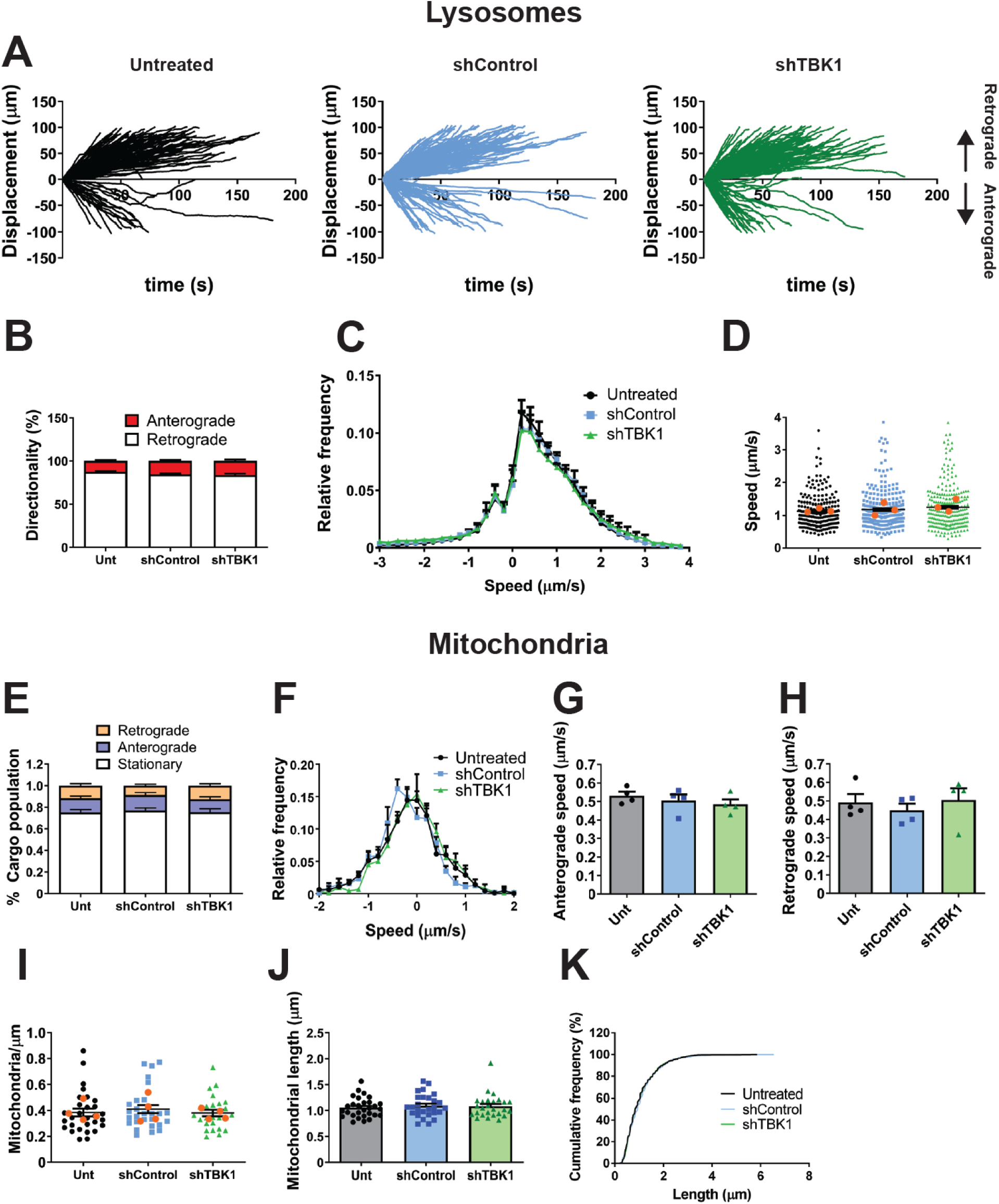
Axonal transport of lysosomes and mitochondria are unaffected by TBK1 knockdown. **(A)** Primary MNs were transduced with either shControl or shTBK1 on DIV 3. 72 h later, MNs were labelled with Lysotracker Deep Red (50 nM) for 30 min, washed and imaged. The displacement graphs for all conditions show bidirectional movement of cargoes, with a bias toward the retrograde direction. **(B)** Quantification of the directionality of transport. Around 80% of lysotracker-positive organelles move retrogradely in all conditions. n = 3; two-way ANOVA with Sidak’s multiple comparison test. **(C)** Speed graphs for every condition, showing overlapping profiles. **(D)** Average speed per cargo, considering every displacement as positive. No statistically significant differences were detected. Number of cargoes: Untreated (Unt) 207, shControl 252, shTBK1 244 from three independent primary cultures, orange dots represent the mean for each replicate; one-way ANOVA with Tukey’s multiple comparison test. **(E)** Primary MNs were transduced with shControl or shTBK1 on DIV 3. 72 h later, MNs were labelled with TMRE (20 nM for 20 min) and imaged. About 75% of mitochondria are static, whilst the rest is split evenly between anterograde and retrograde movement. **(F)** Speed profiles for the three conditions. All profiles overlap almost completely. **(G)** and (**H)** Average anterograde and retrograde speed per experiment, respectively. No statistically significant changes were detected. n = 4, one-way ANOVA with Tukey’s multiple comparison test. **(I)** Mitochondrial density for each group. No significant changes were detected. Number of mitochondria: Unt 29, shControl 27, shTBK1 26 neurons from four independent cultures; one-way ANOVA with Tukey’s multiple comparison test. **(J)** Quantification of mitochondrial length. No significant changes between conditions were detected. Number of mitochondria: Unt 29, shControl 27, shTBK1 26 neurons from four independent cultures; one-way ANOVA with Tukey’s multiple comparison test. **(K)** Mitochondrial length cumulative frequency, showing similar distributions for all conditions.

We then evaluated whether expression of Rab7 WT or the S72 mutants influences the axonal transport of lysosomes. We magnetofected primary MNs in culture with Rab7 WT, S72A or S72E on DIV 5 and tracked lysotracker-labelled endosomes on DIV 6. Mirroring the results obtained with TBK1 knockdown, approximately 80% of lysosomes are transported in the retrograde direction in all conditions (**Supplementary Figure 3A,B),** with overlapping speed profiles (**Supplementary Figure 3C**) and an unchanged average speed per cargo (**Supplementary Figure 3D**).

To further characterise the effects of Rab7 S72 phosphorylation by TBK1 on axonal transport, we decided to monitor the dynamics of a cargo that moves along the axon by a different mechanism. We chose mitochondria, since the recruitment of molecular motors to these organelles depends on the adaptor complex Miro-TRAK1/2 [30], which differs from that of signalling endosomes. We transduced WT primary MNs with shControl or shTBK1 on DIV 3 and tracked mitochondria labelled with tetramethylrhodamine ethyl ester (TMRE) on DIV 6, imaging every 0.5 s. We found that ∼ 75% of mitochondria were static, with the motile fraction equally divided between anterograde and retrograde transport (**Figure 3E**), a distribution that is not affected by TBK1 knockdown. In addition, we detected no difference in the speed profiles between conditions (**Figure 3F**), nor changes in the average speed in the anterograde or retrograde directions (**Figures 3G,H**). We also evaluated possible changes in the axonal distribution of mitochondria (expressed as number of mitochondria per µm of axon length) and mitochondria length, finding that both parameters were identical across conditions (**Figure 3I-K**).

We extended the analysis of mitochondrial axonal transport to MNs expressing Rab7 WT or the S72 mutants. As before, we magnetofected primary MNs with Rab7 WT, S72A or S72E on DIV 5, and tracked mitochondria labelled with TMRE on DIV 6. In agreement with the results shown in **Figure 3**, ∼ 75% of organelles are static, and the fraction of moving mitochondria is equally split between anterograde and retrograde motion (**Supplementary Figure 3E**). This distribution was not changed by the expression of Rab7 WT or the two mutants. The speed profiles overlap for all conditions (**Supplementary Figure 3F**), suggesting that Rab7 WT and the S72 mutants do not alter the axonal transport of mitochondria. In agreement with this, the average speed per cargo in the anterograde and retrograde directions were not significantly different (**Supplementary Figures 3G** and **3H**, respectively). Mitochondrial distribution remained unchanged across conditions (**Supplementary Figure 3I,J**). However, in stark contrast to TBK1 knockdown, we detected an increased proportion of elongated mitochondria upon Rab7 WT and S72A mutant expression (arrows in **Supplementary Figure 3I**, quantified in **Supplementary Figure 3K**). The cumulative frequency of mitochondrial lengths (**Supplementary Figure 3L**) confirms this increased frequency of elongated mitochondria. Altogether, these results rule out the change in axonal transport directionality caused by the loss of TBK1 activity as a global phenomenon, since the dynamics of lysosomes and mitochondria remain largely unaffected.

### Pharmacological modulation of TBK1 activity is cell-type specific

To further characterise the role of TBK1 in the axonal transport of signalling endosomes, we decided to assess how the activation of this kinase might affect their trafficking. First, we checked whether TBK1 activation induced by polyinosinic:polycytidylic acid (p(I:C))-mediated TLR3 stimulation is conserved in MNs. To this end, we treated primary MNs (DIV 6) with either 10 µg/ml or 100 µg/ml p(I:C) for 2 h and measured TBK1 p-S172 levels by western blotting. Despite being widely used to trigger TLR3 activation and TBK1 phosphorylation in a variety of cellular systems [31], p(I:C) was ineffective at increasing the phosphorylation of TBK1 at S172 in primary MNs (**Figure 4A,B**). We also tested whether the TBK1/IKKε inhibitor MRT67307 can be effectively used in our cell model. We pre-treated primary MNs with 2 µM MRT67307 on DIV 6 and stimulated one experimental group with 100 µg/ml p(I:C) after 1 h, treating instead the other group with vehicle. Surprisingly, cells treated with MRT67307 show a trend towards increased TBK1 activation, which is replicated in N2a cells (**Figure 4A,B**). p(I:C) did not modify the effect exerted by MRT67307 on TBK1 p-S172 (**Figure 4A,B**). We also assessed whether p(I:C) was able to activate TBK1 by immunofluorescence. We stimulated primary MNs on DIV 6 with 100 µg/ml p(I:C) for 2 h and then stained them for TBK1 p-S172. As shown in **Figure 4C**, p(I:C) treatment does not increase TBK1 activation. We detected however, a marked accumulation of TBK1 p-S172 near centrosomes in glial cells undergoing cell division (**Figure 4C**), in agreement with previous reports [32]. Since treatment with 10 µg/ml p(I:C) activates TBK1 in Jurkat cells after 120 min (**Supplementary Figure 4A,B**), its inability to do it in MNs indicates the pathway is inactive in this cell type.

**Figure 4.**
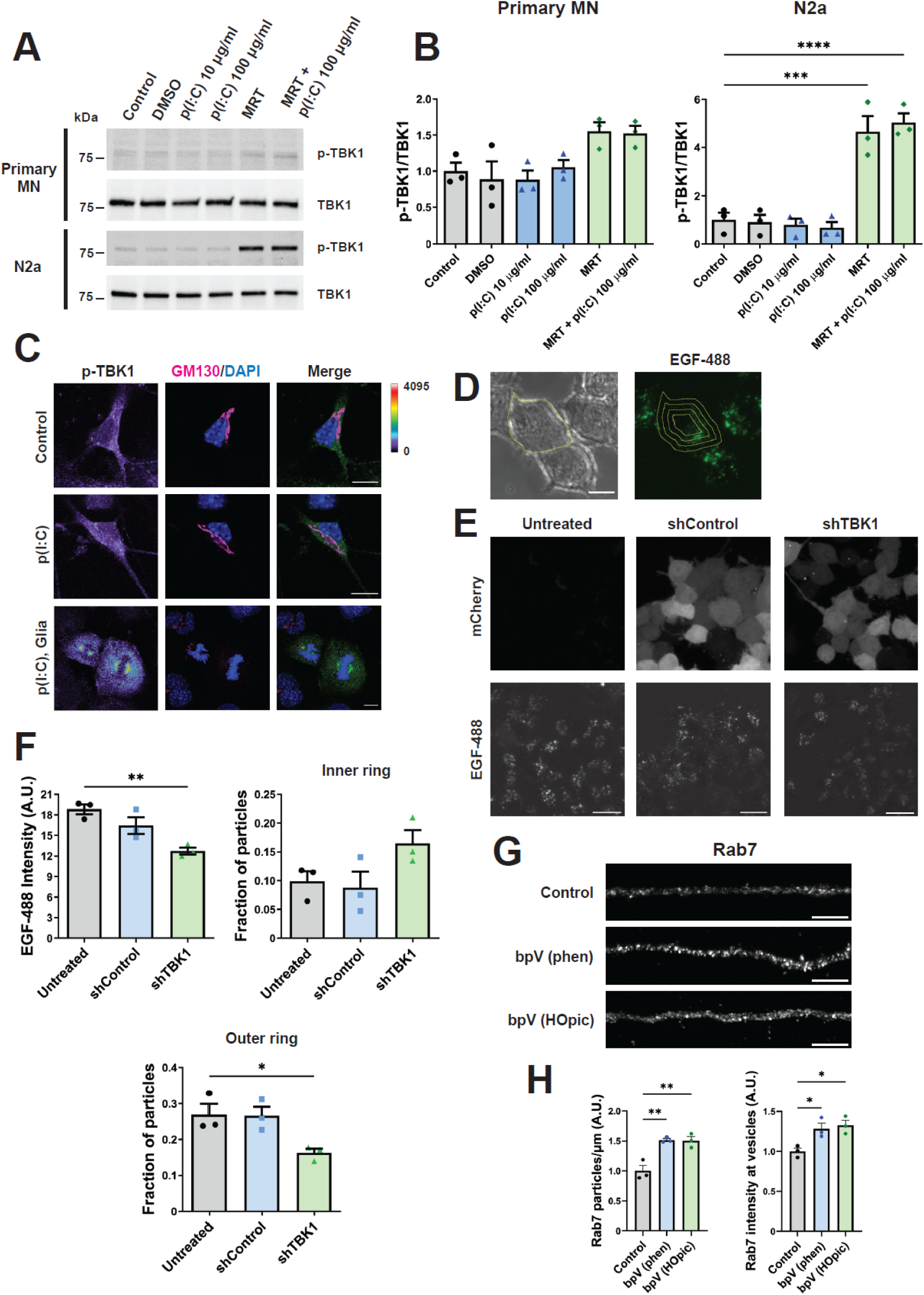
Pharmacological modulation of TBK1 activity is cell-type specific. **(A)** Primary MNs (upper panels) were stimulated for 2 h with 10 μg/ml or 100 μg/ml p(I:C) on DIV 6. Another group was treated with 2 μM MRT67307 for 3 h, or 2 μm MRT67307 for 1 h and 100 μg/ml p(I:C) for another 2 h in the presence of the inhibitor. TBK1 p-S172 and total TBK1 levels were determined by WB. The same conditions were used to stimulate undifferentiated N2a cells (bottom panels). p(I:C) fails to activate TBK1, whilst the TBK1 inhibitor MRT67307 enhances the levels of active TBK1. **(B)** Quantification of TBK1 p-S172/TBK1 from primary MNs (left panel) and N2a cells (right panel) from (A). n = 3, one-way ANOVA with Tukey’s multiple comparison test. ***, P < 0.001, ****, P < 0.0001. **(C)** Primary MNs were stimulated with 100 μg/ml p(I:C) for 2 h on DIV 6, fixed and stained as indicated. TBK1 p-S172 fluorescent intensity levels are shown with a rainbow scale. p(I:C) did not change TBK1 p-S172 levels or its subcellular distribution. An example of non-neuronal cells undergoing mitosis in the same primary culture is presented at the bottom. Centrosomes show clear accumulation of TBK1 p-S172. Scale bar, 10 μm. **(D)** N2a cells were transduced with shControl or shTBK1, stimulated 72 h later with 100 ng/ml EGF-488 for 20 min, and imaged. The cell outline was drawn and eroded inwards 2 µm three times, creating four concentric rings. Scale bar, 10 μm. **(E)** Representative images of EGF-488 uptake, with mCherry expression as the transduction reporter. Scale bar, 20 μm. **(F)** EGF-488 intensity quantification across the whole cell, as well as fraction of particles in the inner or outer ring. TBK1 knockdown decreases the overall EGF signal in N2a cells, whilst also reducing the number of EGF-positive organelles in the cell periphery. n = 3, one-way ANOVA with Tukey’s multiple comparison test, *, P < 0.05, **, P < 0.01. **(G)** Primary MNs were treated on DIV 6 with 100 nM bpV (phen) or bpV (HOpic) for 1 h, followed by fixation and endogenous Rab7 immunolabelling. Scale bar, 5 μm. **(H)** Quantification of the number of Rab7 punta/μm, as well as Rab7 intensity in those puncta. PTEN inhibition increases both Rab7 organelle density, and Rab7 vesicular fluorescent intensity. n = 3, one-way ANOVA with Tukey’s multiple comparison test, *, P < 0.05, **, P < 0.01.

Given that Rab7 p-S72 is dephosphorylated by Phosphatase And Tensin Homolog (PTEN), and this in turn regulates the lysosomal targeting and degradation of epidermal growth factor receptor (EGFR) [15], we tested whether TBK1 knockdown could also affect EGFR trafficking. N2a cells were transduced with shTBK1 or shControl, and 72 h later, stimulated with 100 ng/ml EGF-488 for 20 min and then imaged. We selected the cell outline and then moved it inwards 2 µm three times, creating four cell profiles [33], as shown in **Figure 4D**. TBK1 knockdown reduced EGF-488 overall intensity and significantly decreased the number of EGF-positive organelles located in the most outer ring (**Figure 4E,F**), suggesting that in addition to neurotrophins, other growth factor signalling pathways are affected by TBK1 knockdown. We also tested how PTEN inhibition may impact on Rab7 localisation in WT MNs, and found that two different inhibitors (bpV (phen) and bpV (HOpic)) increase both the number of Rab7-positive organelles and their Rab7 intensity (**Figure 4G,H**), suggesting S72 phosphorylation contributes to the membrane targeting and localisation of Rab7.

### Rab7 phosphorylation by TBK1 at S72 controls recruitment of dynein adaptor RILP

Having shown that TBK1 knockdown induces the bidirectional movement of Rab7-positive signalling endosomes, we sought to identify the mechanism responsible of this phenomenon. First, we purified and tested an antibody against Rab7 p-S72 (**Supplementary Figure 4C,D**). Staining with this antibody yielded a punctate distribution in primary MNs, which as expected, is not changed by p(I:C) stimulation (**Figure 5A**). Of note, cultured MNs triple-stained for endogenous Rab7, TBK1, and H_C_T show that 14.2% ± 0.8% (mean ± SEM, n=3) of H_C_T-positive endosomes also carry TBK1 and Rab7 (**Figure 5B**, white arrows). Interestingly, we also observed Rab7/TBK1 particles without H_C_T, and H_C_T/TBK1 particles without Rab7 (**Figure 5B**, blue and yellow arrows, respectively). We carried on confirming TBK1 phosphorylation on Rab7 S72 by an *in vitro* kinase assay using a library of purified kinases (**Supplementary Table 1**) and Rab7 as substrate. To assess the specificity of the phosphorylation, we also used Rab7 S72P, a loss-of-function mutant that results in embryonic lethality in homozygous mice (data not shown). Whilst several members of the PKC family phosphorylated both Rab7 WT and the S72P mutant, we found that among the kinases tested, only TBK1 and IKKε modify Rab7 uniquely at position S72.

**Figure 5.**
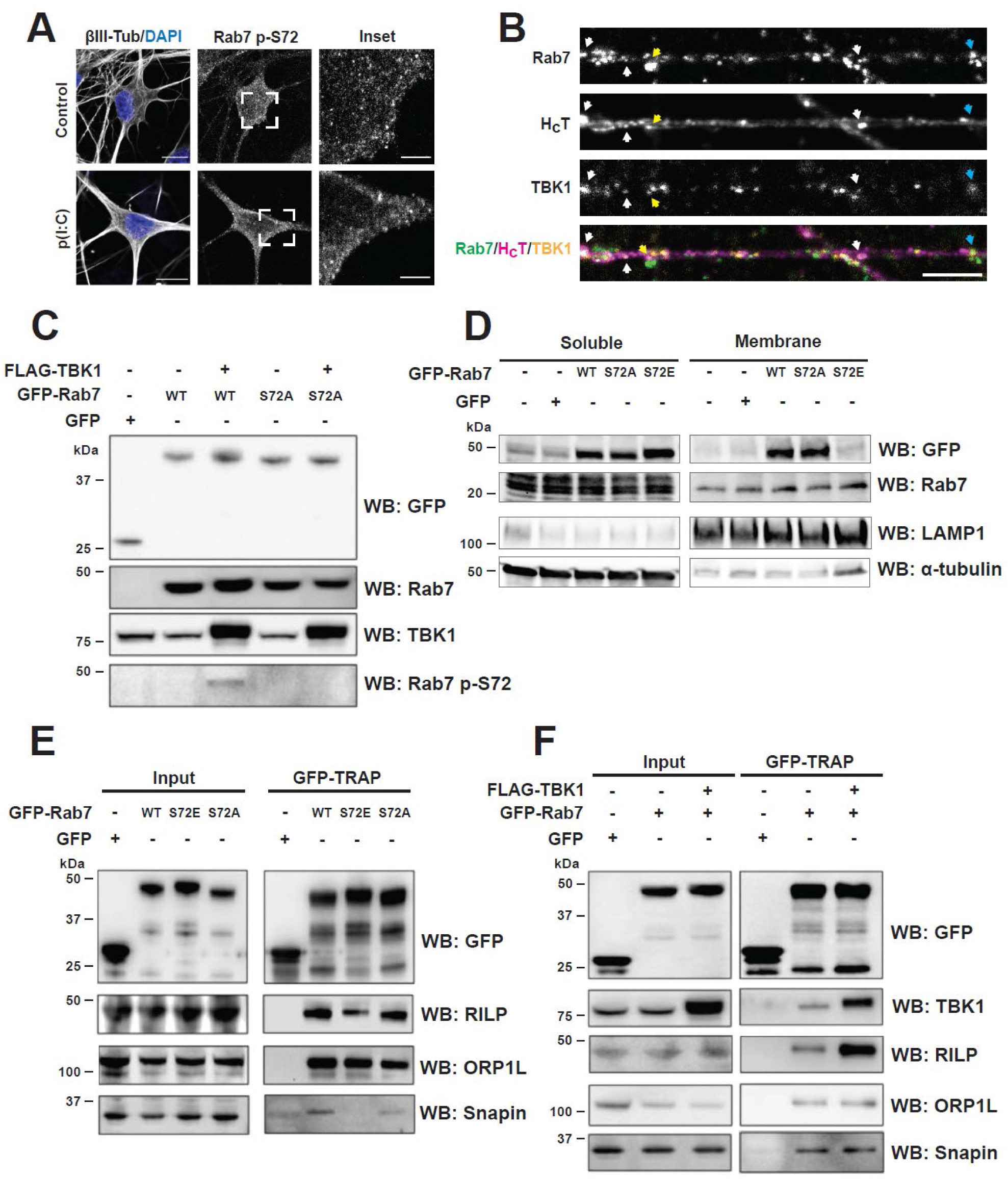
TBK1-dependent Rab7 p-S72 regulates binding to the dynein adaptor RILP. Primary MNs were treated on DIV 6 with 100 μg/ml p(I:C) for 2 h, fixed and stained as indicated. p(I:C) did not alter Rab7 p-S72 levels or its punctate distribution. Scale bar, 10 μm. Insets show examples of Rab7 p-S72 puncta. Scale bar, 3 μm. **(B)** 6 DIV primary MNs were labelled with 30 nM AlexaFluor 555-H_C_T for 45 min, washed and fixed. Double-staining for endogenous Rab7 and TBK1 allows to detect triple-positive cargoes for Rab7/TBK1/H_C_T (white arrows), as well as Rab7/TBK1 organelles (blue arrow) and TBK1/H_C_T carriers (yellow arrow). Scale bar, 5 μm. **(C)** HEK-293 cells were co-transfected with FLAG-TBK1 and either GFP-Rab7 WT or S72A (GFP alone was used as control). The anti-Rab7 p-S72 antibody recognises a band only in the TBK1/Rab7 WT condition. **(D)** N2a cells were transfected with GFP-Rab7 WT or the S72 mutants (including GFP as a control) for 24 h, after which they were used to obtain membrane protein-enriched fractions. LAMP1 and α-tubulin were chosen to monitor the membrane fraction enrichment. Endogenous Rab7 levels in the membrane fraction are not affected by expression of GFP-Rab7 WT, nor by the S72 mutants. **(E)** GFP-TRAP from HEK-293 cells transfected with GFP-Rab7 WT, S72A or S72E as indicated (GFP alone as control). Rab7 S72E binding to both RILP and snapin is decreased, compared to Rab7 WT and S72A. **(F)** GFP-TRAP from HEK-293 cells transfected with FLAG-TBK1 and GFP-Rab7 WT as indicated (GFP alone as control). WB shows RILP, ORP1L and snapin co-precipitation with Rab7. TBK1 expression increases Rab7/RILP interaction.

Next, we checked whether TBK1 overexpression can increase Rab7 p-S72 levels in HEK-293 cells. We co-transfected FLAG-TBK1 with either GFP-Rab7 WT or S72A and detected a band corresponding to Rab7 p-S72 only in the FLAG-TBK1/GFP-Rab7 WT condition (**Figure 5C**), confirming that TBK1 phosphorylates Rab7 at S72 both *in vitro* and *ex vivo*. In addition, we transfected N2a cells with GFP-Rab7 WT, S72A or S72E (using GFP alone as control), and isolated the soluble and membrane fractions, using LAMP1 and α-tubulin enrichment to confirm the fractionation (**Figure 5D**). Our results show that the levels of endogenous Rab7 associated with membranes were not changed by the expression of Rab7 WT or either S72 mutants.

Rab7 phosphorylation at S72 alters its binding to effectors [10,11,15,16]. Hence, we carried out pull-down experiments using recombinant GST-Rab7 either unphosphorylated or phosphorylated *in vitro* by TBK1, to identify interactors whose binding was regulated by Rab7 S72 phosphorylation. Phosphorylated or unphosphorylated Rab7 were incubated with protein lysates from mouse embryonic spinal cord or adult cerebral cortex, followed by GST pull down and quantitative mass spectrometry. **Supplementary Figure 4E** and **F** shows the volcano plots for the differential interactors in embryonic spinal cord and adult cerebral cortex, respectively, and **Supplementary Table 2** and **3** report the list of interactors. We found a trend for dynein heavy chain (Dync1h1) to preferentially interact with Rab7 p-S72, although it was not detected in one of the replicates, hence it did not reach statistical significance.

Interestingly, TBK1 is enriched in the Rab7 p-S72 pull down for both tissues, suggesting this phosphorylation might help recruiting additional TBK1 molecules. We sought to confirm the enhanced dynein binding for Rab7 p-S72 by testing whether dynein adaptors also show the same binding pattern. To this end, HEK-293 cells transfected with GFP-Rab7 WT or the S72 mutants were used for GFP-TRAP immunoprecipitations, followed by immunoblotting for Rab-interacting lysosomal protein (RILP) and oxysterol-binding protein-related protein 1L (ORP1L), two established Rab7 effectors that recruit dynein-dynactin to late endosomes and lysosomes [34,35], as well as snapin, an adaptor that recruits dynein to TrkB signalling endosomes, mediating their retrograde transport in cortical neurons [36]. We found that expression of Rab7 S72E reduced binding to both RILP and snapin, without affecting ORP1L (**Figure 5E**). When the GFP-TRAP experiment was performed with lysates prepared from cells overexpressing FLAG-TBK1 and GFP-Rab7, we found that, in contrast to the Rab7 S72E condition, Rab7 S72 phosphorylation increases its interaction with RILP, in agreement with the interactomics results, whilst binding to ORP1L and snapin was unaffected (**Figure 5F**). This discrepancy between the S72E mutant and TBK1-phosphorylated Rab7 likely arises from the Rab7 S72E cytosolic mis-localisation and consequent loss-of-function [10,15].

Altogether, these results show TBK1 ensures the unidirectional transport of neurotrophic signalling endosomes, through Rab7 S72 phosphorylation and the recruitment of cytoplasmic dynein. Since the axonal transport of other organelles was not affected in our study, this suggests compartment-specific mechanism(s) of motor recruitment modulated by TBK1 in MNs.

## Discussion

Phosphorylation has emerged as a main regulatory mechanism for Rab GTPases, with profound implications for intracellular trafficking deregulation during disease. Here, we report that TBK1-dependent Rab7 p-S72 maintains the unidirectional transport of neurotrophic signalling endosomes in MNs, through the recruitment of the dynein adaptor RILP. Interestingly, Rab7 S72E overexpression had a similar effect to TBK1 knockdown. Since this mutant shows reduced interaction with components of the geranylgeranyltransferase complex, as well as with GDP dissociation inhibitors [10,15], it is predicted to act as a loss-of-function, rather than a phospho-mimetic, mutant. Even though Rab7 S72E does not cycle through membrane compartments, we decided to use it as a discovery tool since this mutant has been successfully used to identify new Rab7 p-S72 interactors, such as folliculin [10]. Our results show that whilst expression of the dominant-negative mutant Rab7 N125I blocks signalling endosome transport [4], Rab7 S72E only affects their direction of transport, hence it is likely to disrupt only Rab7 p-S72-dependent processes.

TBK1 regulates axonal transport directionality in a manner that is specific for signalling endosomes, as lysosomes remain unaffected upon TBK1 downregulation. A similar specificity has been previously observed in signalling endosomes carrying TrkB in hippocampal neurons, whose trafficking relies on the dynein adaptor Hook1 [37]. In this model, Hook1 knockdown reduces signalling endosome processivity without affecting the dynamics of LC3B-positive autophagosomes or mitochondria.

A possible explanation for the lack of a global effect of TBK1 on transport might be that this kinase is only present in a sub-population of Rab7 organelles. For instance, a recent report showed occasional TBK1 association with lysosomes in HeLa cells, which increases after TBK1 overexpression [21]. Similarly, we found that only 15% of signalling endosomes carry both Rab7 and TBK1 in MNs, pointing towards a fine regulation of TBK1 localisation.

The axonal transport of mitochondria was also unaltered by TBK1 knockdown. This result is in agreement with a study analysing ALS-linked TBK1 mutations impairing dimer formation and/or catalytic activity, which found no change in somatic mitochondrial mass upon expression of these mutants in hippocampal neurons in basal conditions [18]. However, we cannot rule out a possible effect of TBK1 knockdown on mitochondrial homeostasis if mitochondrial function is challenged by depolarising agents. Alternatively, the residual levels of TBK1 after treatment with shRNAs might still be sufficient to ensure some of its functions. We detected, however, an increase in mitochondrial length in MNs expressing Rab7 S72A. Given the established role of Rab7 in mitochondrial fission at mitochondrion-lysosome contact sites [38], our results suggest this function might also be modulated by TBK1-dependent phosphorylation of Rab7. Altogether, these observations indicate the regulation of the axonal transport of signalling endosomes is highly specific.

The bidirectional movement of signalling endosomes after TBK1 knockdown points towards an imbalance in the recruitment of molecular motors, which may engage in a “tug-of-war” [39]. To gain mechanistic insights on this process, we looked for components of the cytoplasmic dynein complex, kinesin family members, or their adaptors, showing a preferential binding for Rab7 p-S72, using lysates from embryonic spinal cord and adult brain cortex. However, we were unable to detect motor adaptors among the Rab7 interactors under our experimental conditions. Similarly, motor adaptor proteins were not detected in a proximity-labelling study of autophagosomes in TBK1 E696K/E696K mouse embryonic fibroblasts [40]. These negative results suggest that the experimental conditions to preserve the interactome of motor adaptors and their reliable mass spec detection require further optimisation. Nevertheless, we detected an enrichment in cytoplasmic dynein heavy chain in the Rab7 p-S72 pull down, which, although did not reach statistical significance, prompted us to further investigate the binding of dynein adaptors by biochemical approaches.

RILP is one of the best characterised Rab7-binding proteins involved in the recruitment of cytoplasmic dynein to lysosomes [34,35]. However, the effect of Rab7 phosphorylation on RILP binding remains controversial. Conflicting reports have found phosphorylation at S72 increases, decreases or does not change the strength of Rab7 binding to RILP [11,15,41]. In our experimental conditions, we found that Rab7 p-S72 promotes binding to RILP; whilst the S72E mutants shows lower affinity to RILP and negligible binding to snapin, further supporting its loss-of-function characteristics. In contrast, Rab7 S72A still interacts with RILP, in agreement with previous reports [15]. This indicates that S72 phosphorylation, rather than being required for RILP binding, has a modulatory role, enhancing or stabilising this association. Henceforth, we propose that TBK1 ensures the unidirectional transport of signalling endosomes by enhancing the recruitment of the retrograde motor dynein through its adaptor RILP to the surface of these organelles.

Surprisingly, the machinery for anterograde movement is also associated to signalling endosomes, since the proteome of H_C_T-positive organelles isolated from mouse embryonic stem cell-derived MNs contains the kinesin-1 heavy chains KIF5B and KIF5C, the kinesin light chain 1 and 2, and Arl8b [42], a small GTPase having kinesin 1 and 3 as its effectors [43]. In this light, the Rab7 p-S72 switch may function as a toggle mechanism allowing the switch between motors with different directionality.

The modulation of signalling endosome transport directionality exerted by TBK1, presents similarities with the mechanisms by which LRRK2 controls the trafficking of LC3-positive autophagosomes [44], albeit with significant differences. For instance, the axonal transport of autophagosomes is unidirectional in basal conditions but becomes bidirectional upon expression of the overactive LRRK2 G2019S mutant. In contrast, signalling endosomes move bidirectionally upon TBK1 downregulation. Furthermore, the retrograde movement of signalling endosomes relies on Rab7, whilst in autophagosomes, LRRK2 acts through a mechanism involving Rab10, ARF6 and the dynein adaptors JIP3 and JIP4 [39,45]. Upon LRRK2 phosphorylation, p-Rab8 and p-Rab10 are able to recruit additional LRRK2 to these organelles, which in turn enhances the kinase activity, establishing a feed-forward mechanism [46]. Similarly, our mass spec analysis showed that TBK1 was highly enriched in the Rab7 p-S72 sample. Although we cannot rule out that this result might be partially due to the *in vitro* kinase approach used to generate Rab7 p-S72, it is tempting to speculate the existence of a TBK1/Rab7 p-S72 feed-forward mechanism allowing for the recruitment of additional TBK1 onto signalling endosomes.

Since TBK1 loss-of-function alters the axonal transport of signalling endosomes, we attempted to test whether TBK1 activation also has an effect. Unfortunately, TBK1 overexpression was toxic for MNs, whilst established TBK1 pharmacological activators, such as p(I:C), were ineffective in MNs and N2a cells. The latter result is likely to be due to the low expression of Toll-like receptor 3 (TLR3), the receptor that recognises p(I:C), in MNs. Despite TLR3 being one of the highest TLRs expressed in this neuronal subtype, its levels are still an order of magnitude lower than in peripheral macrophages [47].

Rab7 S72 is also differentially phosphorylated in a cell type-dependent manner. For example, p-S72 can be induced by phorbol 12-myristate 13-acetate (PMA) in mouse embryonic fibroblasts, but not in the human skin keratinocyte cell line HaCaT [41]. Conversely, HaCaT cells show increased Rab7 p-S72 levels after treatment with EGF [41], highlighting the importance of studying this signalling cascade in the cell type of interest. Unexpectedly, we found that the established TBK1 inhibitor MRT67307 activates TBK1 in N2a cells and to a lesser extent in MNs. Other studies have shown that the TBK1 inhibitors BX795, MRT67307 and GSK8612 do not completely prevent TBK1 trans-autophosphorylation at p-S172, hinting at the existence of upstream kinases activating TBK1 [48].

Given the deficits in axonal transport of signalling endosomes observed in ALS models [6,7], what could be the contribution of TBK1 loss-of-function to disease pathogenesis? TBK1 deficiency has been linked to endolysosomal dysfunction. *TBK1*^-/-^ human MNs exhibit impaired endosomal maturation and reduced lysosomal function, which results in TAR DNA-binding protein 43 (TDP-43) mislocalisation to the cytoplasm [49]. This effect can also be triggered by poly(GA) derived from C9orf72 (G_4_C_2_) expansion, with poly(GA) inclusions sequestering TBK1, which in turn induces early endosome enlargement and TDP-43 aggregation, a phenotype that is worsened by the ALS-linked TBK1 R228H mutation [22]. In addition, mice homozygous for the ALS-causing TBK1 E696K mutation show an enrichment of enlarged lysosomes lacking degradative enzymes in spinal cord MNs, pointing towards a disrupted lysosomal biogenesis and/or turnover [40]. Thus, the abnormal bidirectional transport of signalling endosomes could be another manifestation of how this pathway is affected by TBK1 loss-of-function, specifically in MNs.

TBK1 exhibits a biphasic function during ALS pathology, being protective at early stages in a cell-autonomous manner, but detrimental at later stages, mainly due to non-cell autonomous mechanisms related to its role in inflammation [25]. We hypothesise that reduced TBK1 activity compromises the delivery of pro-survival signals carried by signalling endosomes, impacting long-term motor neuron survival, and fitting with the protective role of TBK1 at disease onset. As the reversion in transport is partial, it is possible that the negative effect on survival only appears at a late stage, or it requires a “second hit”, in line with recent TBK1 ALS models exhibiting mild but steady accumulation of subtle cellular phenotypes [40].

In summary, we report a novel neuronal role for TBK1 in controlling the directionality of axonal transport of neurotrophic signalling endosomes through the phosphorylation of Rab7 at S72 and the recruitment of the dynein adaptor RILP. This effect is specific for these organelles, highlighting a new mechanism for differential axonal transport regulation of distinct cargoes in MNs.

## Material and Methods

### Animals

The Ethical Approval is informed in the Declarations section. Wild type mice from a C57BL/6 x SJL background were used in all experiments.

### Antibodies and reagents

In this work, we used the following antibodies: anti-acetylated tubulin (Santa Cruz, sc-23950, 1:400), anti-α tubulin (Abcam, ab6161, 1:2,000), anti-βIII tubulin (Tuj1) (Covance, MMS-435P, 1:1,000), anti-GM130 (BD Transduction laboratories, #610822, 1:100), anti-GFP (Santa Cruz, sc-9996, 1:2,500), anti-LAMP1 (Santa Cruz, sc-5570, 1:1,000), anti-MAP2 (Chemicon, AB5622, 1:1,000), anti-ORP1L (R247, a gift from Vesa Olkkonen, 1:1,000), anti-TBK1 p-S172 (Cell Signaling, mAb#5483, 1:500 in western blot (WB), 1:200 in immunofluorescence (IF)), anti-Rab7 (Abcam, ab50533, 1:1,000 in WB, 1:250 in IF), anti-RFP (GenScript, A00682, 1:1,000), anti-RILP (Abcam, ab140188, 1:1,000), anti-SMI-31 (Covance, SMI-31R, 1:1,000), anti-snapin (Synaptic Systems, 148 002, 1:3,000), anti-TBK1 (Abcam, ab40676, 1:3,000 in WB, 1:200 in IF) and anti-tyrosinated tubulin (Millipore, mab1864, 1:500). Lysotracker Red DND-99, lysotracker Deep Red and TMRE (T669; 1 mM stock) were purchased from ThermoFisher. Poly(I:C) LMW (p(I:C)) and EGF-488 were from Invitrogen, and bpV (phen), bpV (HOpic) and MRT67307 were obtained from Sigma.

### Anti Rab7 p-S72 antibody generation and purification

A peptide comprising the sequence flanking Rab7 S72 was synthesised with S72 phosphorylated (CERFQpSLGVA-CONH_2_) and used for the immunisation of two rabbits by BioGenes GmbH (Berlin, Germany). For antibody purification, phosphorylated and unphosphorylated forms of the peptide were dissolved in coupling buffer (50 mM Tris-HCl pH 8.5, 5 mM EDTA), added to Sulfolink resin (ThermoFisher) and incubated for 1 h. Excess peptide was removed and the reaction stopped by adding coupling buffer containing 50 mM cysteine (pH 8.0) for 30 min. Resins were packed into separated columns and washed sequentially with water, wash buffer 2 (1 M NaCl, 200 mM glycine pH 2.4) and water at 4°C. The hyperimmune serum was diluted 1:1 in Tris-buffered saline (TBS: 50 mM Tris-HCl, pH 7.4, 150 mM NaCl), added to the unphosphorylated peptide column and incubated at 4°C for 2 h under rotation. The flow through was then added to the phosphorylated peptide column and incubated at 4°C for 2 h under rotation. The column was washed twice with ice-cold TBS, and the antibody eluted in 200 mM glycine, pH 2.4. The eluate was immediately neutralized with 1.5 M Tris-HCl, pH 8.8 and bovine serum albumin (BSA) was added to 0.1% final concentration. After purification, the antibody was tested by ELISA, as follows: phosphorylated and unphosphorylated peptides were diluted in TBS, 0.01 – 10 ng added to a 96-wells plate and allowed to air-dry overnight. The next day, wells were blocked with TBS containing 0.1 % Tween-20 and 1% BSA for 1 h at room temperature. The purified antibody was incubated for 2 h, wells washed three times, and an anti-rabbit secondary antibody conjugated to horseradish peroxidase (HRP) was added for 1 h. Wells were then washed five times before addition of SIGMAFAST OPD substrate (P9187, Sigma) for 10 min. The reaction was stopped by adding 50 µl 2 M HCl and the absorbance was measured at 450 nm.

### Plasmids

Canine Rab7 WT cloned into the mammalian expression vector pEGFF-C1 was kindly provided by Cecilia Bucci [50]. Rab7 S72A and S72E were generated by site-directed mutagenesis (QuikChange Site-Directed Mutagenesis Kit, Stratagene). EB3-mCherry was a gift from Michael Davidson (Addgene plasmid # 55037). A shRNA set directed against TBK1 plus a control sequence were purchased from GeneCopoeia (# MSH101526-LVRU6MP), with an U6 promoter and mCherry as reporter gene. Control sequence: GCTTCGCGCCGTAGTCTTA, shRNA 1: CCAGAATCAGAATTTCTCATT, shRNA 2: CCAGTTCTTGCAAACATACTT.

### *In vitro* kinase assays

The *in vitro* kinase validation was carried out by ProQinase GmbH, using a radiometric protein kinase filter-binding assay and a library of 190 serine/threonine kinases, using His_6_-Rab7 WT or S72P as substrates. *In vitro* phosphorylation of GST-Rab7 by TBK1 (ThermoFisher) was performed in reaction buffer (20 mM Tris-HCl pH 7.5, 10 mM MgCl_2_, 1 mM EGTA, 10 µM Na_3_VO_4_, 0.5 mM β-glycerophosphate, 2 mM dithiothreitol (DTT) and 0.01% Triton X-100) with 100 µM ATP and 100 ng of TBK1 per 50 µl of reaction. Samples were incubated at 30°C for 30-60 min, boiled and analysed by western blotting.

### Cell culture

Tissue culture reagents were purchased from ThermoFisher, unless stated otherwise. N2a and HEK cells were maintained in DMEM supplemented with 10% FBS and 1% GlutaMAX at 37°C and 5% CO_2_. Jurkat cells were grown in RPMI media with 10% FBS at 37°C and 5% CO_2_. Cells were routinely tested for mycoplasma using a MycoAlert detection kit (Lonza). Primary MNs were isolated as previously described [26,51]. MNs were resuspended in motor neuron medium (Neurobasal, 2% heat-inactivated horse serum, 2% B27 supplement, 1X GlutaMax, 25 μM 2-mercaptoethanol, 1% penicillin/streptomycin, recombinant rat CNTF (R&D Systems, 10 ng/ml), recombinant rat GDNF (R&D Systems, 0.1 ng/ml) and recombinant human BDNF (R&D Systems, 1 ng/ml)). Magnetofection (NeuroMag, OZ Biosciences) was carried out following manufacturer’s instructions (with 0.5 µg of DNA, 1.75 µl beads and 15 min of incubation on top of a permanent magnet). Magnetofected neurons were incubated for 18-24 h and used for further experiments.

### Western blotting

For WB, cells were lysed in RIPA buffer (50 mM Tris-HCl, pH 7.5, 1 mM EDTA, 2 mM EGTA, 150 mM NaCl, 1% NP40, 0.5% sodium deoxycholate and 0.1% sodium dodecylsulfate containing protease and phosphatase inhibitors (HALT, ThermoFisher)), for 30 min at 4°C. Lysates were centrifugated at 14,000 rpm for 15 min at 4°C and protein concentration was determined. Between 10 and 20 μg of total protein extract were loaded on 4-15% Mini-PROTEAN TGX pre-cast gels (BioRad) or NuPAGE 4-12% Bis-Tris gels (ThermoFisher) and transferred into polyvinylidene difluoride (PVDF) membranes (BioRad). Membranes were blocked in TBS containing 0.05% Tween-20 and 5% BSA for 1 h at room temperature and then incubated with primary antibodies overnight at 4°C. Membranes were washed and incubated with horseradish peroxidase (HRP)-conjugated secondary antibodies (GE Healthcare) for 1 h at room temperature. Immunoreactivity was detected using Immobilon Classico ECL substrate (Millipore) and a ChemiDoc MP Imaging System (BioRad).

### Membrane fractionation

N2a cells were transfected with GFP-Rab7 WT, S72A or S72E using Lipofectamine 3000 (ThermoFisher) in Opti-MEM (Gibco) at 70% confluency for 24 h. Cells were then used for the purification of soluble/membrane-enriched protein extracts, with the Mem-PER^TM^ Plus Membrane Protein Extraction Kit (ThermoFisher), following manufacturer’s instructions. The fractions enriched in cytosolic or membrane proteins were used to determine Rab7 levels by WB.

### Immunoprecipitation

GFP-TRAP (Chromotek) immunoprecipitations were performed using HEK-293 cells transfected with Lipofectamine 3000 in Opti-MEM at 70% confluency for 24 or 48 h. Cells were lysed in TBS Lysis buffer (10 mM Tris-HCl pH 7.5, 150 mM NaCl, 0.5 % Nonidet™ P40 Substitute) containing HALT protease and phosphatase inhibitors by pipetting and keeping them on ice for 30 min. Cell lysates were then centrifuged for 10 min at 15,000 x *g* at 4°C, the supernatant collected and then incubated with equilibrated GFP-TRAP beads at 4°C for 1 h under rotation. Beads were then washed with TBS three times in a magnetic rack. Bound proteins were eluted from the GFP-TRAP beads using NuPAGE 2X LDS sample buffer (ThermoFisher) with 100 mM DTT (Sigma-Aldrich) at 99°C. Supernatants were separated from the beads using a magnetic rack, and samples were then processed for WB.

GST-TRAP immunoprecipitations were also carried out using protein extracts from mouse embryonic spinal cord (E12.5) and adult brain cortex lysed using NP40 buffer (50 mM Tris-HCl pH 8.0, 150 mM NaCl, 1% NP-40) containing HALT protease and phosphatase inhibitors. After keeping the homogenates for 30 min on ice, lysates were centrifugated at 14,000 rpm for 15 min at 4°C. The supernatants were incubated with GST-TRAP beads previously bound to recombinant GST-Rab7 or TBK1-phosphorylated Rab7 at 4°C, overnight under rotation. Beads were then washed with NP40 buffer three times and kept for additional processing by mass spectrometry.

### Mass spectrometry and validation

GST-TRAP beads bound to phosphorylated or unphosphorylated Rab7 and treated as described in the previous section were reconstituted in 100 mM Tris-HCl (pH 8.5) and 4% sodium deoxycholate. Proteins were subjected to proteolysis with 1:50 trypsin overnight at 37°C with constant shaking. Digestion was stopped by adding 1% trifluoroacetic acid to a final concentration of 0.5%. Precipitated sodium deoxycholate was removed by centrifugation at 10,000 x *g* for 5 min, and the supernatant containing digested peptides was desalted on an SOLAµ HRP (ThermoFisher). Peptides were dried and dissolved in 2% formic acid before liquid chromatography–tandem mass spectrometry (MS/MS) analysis. A total of 2,000 ng of the mixture of tryptic peptides was analysed using an Ultimate3000 high-performance liquid chromatography system coupled online to an Eclipse mass spectrometer (ThermoFisher). Buffer A consisted of water acidified with 0.1% formic acid, whilst buffer B was 80% acetonitrile and 20% water with 0.1% formic acid. The peptides were first trapped for 1 min at 30 μl/min with 100% buffer A on a trap (0.3 mm by 5 mm with PepMap C18, 5 μm, 100 Å; ThermoFisher). The resulting peptides were separated by a 50 cm analytical column (Acclaim PepMap, 3 μm; ThermoFisher). The gradient was 9 to 35% B in 44 min at 300 nl/min. Buffer B was then raised to 55% in 2 min and increased to 99% for the cleaning step. Peptides were ionized using a spray voltage of 2.1 kV and a capillary heated at 280°C. The mass spectrometer was set to acquire full-scan MS spectra (350 to 1,400 mass/charge ratio) for a maximum injection time set to Auto at a mass resolution of 60,000 and an automated gain control (AGC) target value of 100%. For a second the most intense precursor ions were selected for MS/MS. HCD fragmentation was performed in the HCD cell, with the readout in the Orbitrap mass analyser at a resolution of 30,000 (isolation window of 1.4 Th) and an AGC target value of 200% with a maximum injection time set to Auto and a normalized collision energy of 30%.

All raw files were analysed by MaxQuant v2.1 software using the integrated Andromeda search engine and searched against the Mouse UniProt Reference Proteome (55,029 protein sequences). MaxQuant was used with the standard parameters with only the addition of deamidation (N) as variable modification. The experiment was repeated four times, and the three replicates more consistent between them, according to a principal component analysis, were subjected to a second filter, where only interactors detected in a minimum of two replicates in at least one condition were kept. The analysis of differentially enriched interactors was carried out in R, here, protein intensity data were log₂-transformed, and missing values were imputed using a modified approach to preserve statistical power whilst minimizing bias. Zero values were first replaced with 1 to allow log transformation, after which any remaining zeros were substituted with random values sampled from a normal distribution (μ = 2, σ = 1). Differentially expressed proteins were identified using a moderated t-test approach implemented in the limma package (v3.64.0). A linear model was fitted to the log₂-transformed intensities with a design matrix contrasting experimental conditions. Empirical Bayes moderation was applied to stabilize variance estimates, and significance was assessed using false discovery rate (FDR)-adjusted p-values (q < 0.1). Results were filtered to exclude keratins as potential contaminants. Volcano plots were generated with the VolcaNoseR webtool [52].

### Immunofluorescence

Cells were fixed in 4% paraformaldehyde and 4% sucrose in phosphate-buffered saline (PBS) for 30 min at room temperature, washed three times with PBS, permeabilised with Triton X-100 0.2% in PBS for 5 min, washed again three times and blocked in 5% BSA in PBS for 1 h at room temperature. Primary antibodies were incubated overnight at 4°C in blocking solution. Coverslips were washed three times, incubated with AlexaFluor-conjugated secondary antibodies for 1 h at room temperature, washed again three times, stained with 4′,6-diamidino-2-phenylindole (DAPI) when indicated, washed once with water and mounted on Mowiol. TBK1 localisation experiments in MNs were carried out in the same way, except that cells were permeabilised with saponin 0.05% in PBS for 5 min, and blocked with 5% BSA, 0.05% saponin in PBS for 1 h. Primary and secondary antibodies were incubated in the same blocking solution. For the tyrosinated/acetylated tubulin analysis, MNs were magnetofected on DIV 2. After 24 h, neurons were simultaneously extracted and fixed, as previously described [53]. Briefly, MNs were washed with PBS and then extracted/fixed with a solution containing PHEM (60 mM PIPES-NaOH, 25 mM HEPES-NaOH, 10 mM EGTA and 2 mM MgCl_2_, pH 6.9), 4% paraformaldehyde, 0.15% glutaraldehyde and 0.2% Triton X-100 for 15 min. After this, the immunofluorescence protocol was carried out as described above.

### Immunofluorescence quantification

Fluorescence intensity profiles for both tyrosinated and acetylated tubulin were analysed with Fiji ImageJ [54]. Profiles were measured starting at the axonal tip towards the cell body, until obtaining a profile of at least 50 μm. The ratio between both channels on the proximal and distal axon was determined as: average measurement from segments of 5 μm in length, between 5 and 10 μm from the axon tip (shown as distal) and between 45 and 50 μm from the axonal tip (shown as proximal). For the analysis of H_C_T accumulation at neurite tips, we measured the mean fluorescent intensity in the last 10 μm of each distal process.

For the EGF-488 endosomes subcellular distribution experiment, we outlined the border of each cell analysed, and then enlarged the selection by -2 μm (3 times), as reported previously [33], before calculating the fraction of particles in the inner or outer ring.

For the determination of Rab7 fluorescence associated to vesicles in control neurons, or after PTEN inhibition, we created a mask for Rab7-positive particles with a threshold above the Rab7 cytosolic signal, using Fiji ImageJ, followed by counting number of particles and measuring fluorescent intensity in those particles. Similarly, to quantify the number of Rab7/ H_C_T/TBK1 endosomes, we identified H_C_T-positive structures and checked what percentage were also positive for the other two endogenous fluorescent signals.

### Live cell imaging and tracking

Primary MNs were transduced on DIV 3 or magnetofected on DIV 5, as described above. On DIV 6, neurons were labelled with either 30 nM AlexaFluor-conjugated H_C_T for 45 min, 50 nM Lysotracker Red or Deep Red for 30 min, or 20 nM TMRE for 20 min. Cells were washed and imaged in motor neuron medium supplemented with 20 mM HEPES-NaOH pH 7.4 (for mitochondrial transport experiments, the imaging medium also contained 20 nM TMRE). Images were acquired every 0.5 s, for a total of 400 images per time-lapse (300 frames for mitochondria experiments). Endosomes and lysosomes were tracked using the Fiji ImageJ plugin TrackMate [55] in manual mode. Only endosomes moving for at least ten frames were tracked, until they exited the imaging window or reached a terminal pausing, which corresponds to the absence of movement for more than 10 frames. Mitochondria were tracked using the ImageJ plugin KymoAnalyzer [56] with the following parameters: pixel size: 0.1 μm, frame rate: 2 frames/s, multiplication factor in “Segments” plugin (plugin 5): 2, threshold for the detection of mobile/stationary and switching tracks: 0.5 µm and threshold for the assignment of segments/pauses: 0.1. For the EB3 comets experiments, MNs were magnetofected on DIV 2 and imaged after 6 h. Images were acquired every 0.5 s, for a total of 100 images per time-lapse. Kymographs for the signalling endosomes transport and EB3-comets experiments were generated with the Multi Kymographs analysis option in Fiji ImageJ.

### Statistical analysis

Statistical analysis was performed using GraphPad Prism software. Data are shown as mean ± SEM. We compared multiple groups by one-way ANOVA with Tukey’s multiple comparison test and grouped conditions by two-way ANOVA with Sidak’s multiple comparison test. For cases when variances were non-comparable, Kruskal-Wallis non-parametric one-way ANOVA was used. We used D’Agostino-Pearson as a normality test. Statistical significance was considered as follows: *, P < 0.05, **, P < 0.01, ***, P < 0.001 and ****, P < 0.0001.

## Supplementary Material

**Supplementary Figure S1** presents the validation of the TBK1 knockdown and its effect on primary MN survival. **Supplementary Figure S2** rules out polarity defects or changes in endosomal sorting as the cause of altered axonal transport. **Supplementary Figure S3** shows that Rab7 S72E does not impact the axonal transport of lysosomes or mitochondria. **Supplementary Figure S4** demonstrates that TBK1 activation by p(I:C) is cell-type specific. In addition, it presents the validation of our Rab7 p-S72 antibody and reports the interactors which differentially bind to Rab7 p-S72. **Supplementary Table 1** reports the kinases tested in the Ser/Thr kinase screen. **Supplementary Table 2** lists the Rab7 p-S72 interactors found in embryonic spinal cord, with their respective fold-change versus unphosphorylated Rab7 and their p-values. **Supplementary Table 3** lists Rab7 p-S72 interactors found in adult cortex, including their fold-change versus unphosphorylated Rab7 and their p-values.

## Declarations

### Ethics approval and consent to participate

All experiments were carried out following the guidelines of the Queen Square Institute of Neurology Genetic Manipulation and Ethic Committees, and in accordance with the European Community Council Directive of November 24, 1986 (86/609/EEC). Animal experiments were undertaken under license from the UK Home Office in accordance with the Animals (Scientific Procedures) Act 1986 (Amended Regulations 2012).

### Consent for publication

Not applicable.

### Author contributions

D. Villarroel-Campos: conceptualisation, formal analysis, investigation, project administration, resources, validation, visualisation, and writing (original draft). J.N. Vargas: conceptualisation, formal analysis, investigation, validation, and visualisation. R. Zenezini Chiozzi: formal analysis and investigation. A-L. Brown: formal analysis. M. Wallace: formal analysis and investigation. K. Sun: investigation. J.N. Sleigh: formal analysis, funding acquisition and investigation. K. Thalassinos: Supervision. P. Fratta: conceptualisation and funding acquisition. G. Schiavo: conceptualisation, funding acquisition, project administration, resources, supervision, and writing (original draft). All authors approved the final version of this manuscript and submission of this work.

## Acknowledgements

We thank members of the Molecular NeuroPathobiology (MNP) and Greensmith laboratories (UCL Queen Square Institute of Neurology) for critical reading of the manuscript and personnel of the Denny Brown Laboratory (UCL) for assistance with mouse colonies. All data required to evaluate the conclusions in the paper are present in the paper and/or in the Supplementary Materials.

## Supplementary Figures legends

**Supplementary Figure 1.**
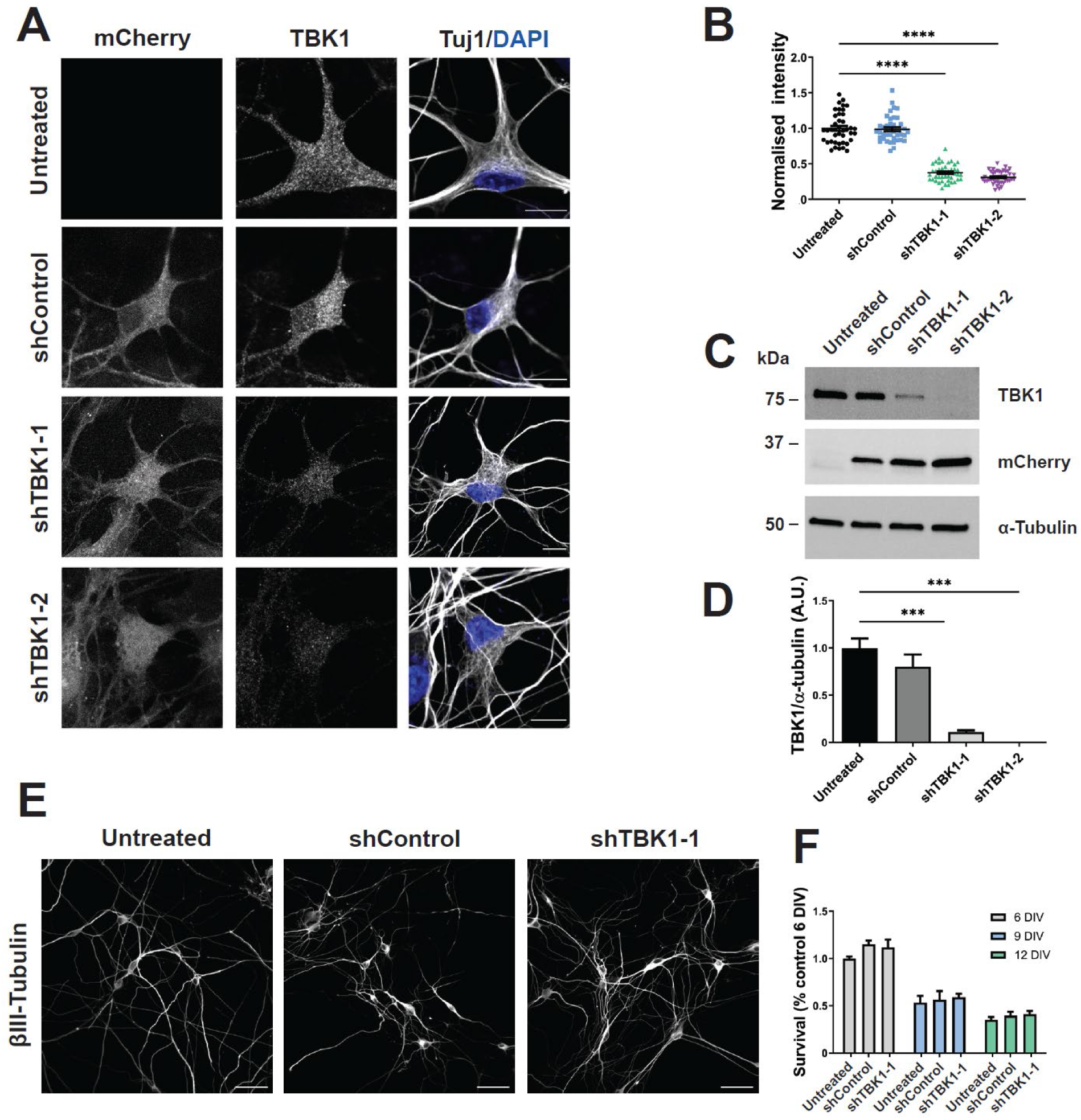
TBK1 knockdown validation. **(A)** Primary MNs were transduced with shControl, shTBK1-1 or shTBK1-2 on DIV 3. Neurons were fixed 72 h later and stained as indicated. Scale bar, 10 μm. **(B)** Fluorescence intensity quantification from transduced MNs in (A). We observed a knockdown efficiency of ∼ 65% for both shRNAs. Number of neurons: Untreated 41, shControl 40, shTBK1-1 43, shTBK1-2 44, from three independent cultures; one-way ANOVA with Tukey’s multiple comparison test, ****, P<0.0001. **(C)** Primary MNs were transduced as in (A), and TBK1 expression levels were determined by WB. **(D)** TBK1 knockdown quantification from (C), normalised to untreated levels. We obtained ∼ 90% knockdown with shTBK1-1 and almost complete silencing with shTBK1-2, whilst shControl was not significantly different from untreated MNs. n = 3; one-way ANOVA with Tukey’s multiple comparison test, ***, P<0.001. **(E)** Primary MNs were transduced with shControl or shTBK1-1 on DIV 3, and fixed on DIV 6, 9 and 12. Scale bar, 50 μm. **(F)** MN survival quantification after TBK1 knockdown, compared to control neurons on DIV 6. TBK1 knockdown does not alter MN viability. Two-way ANOVA with Šidak’s multiple comparison test.

**Supplementary Figure 2.**
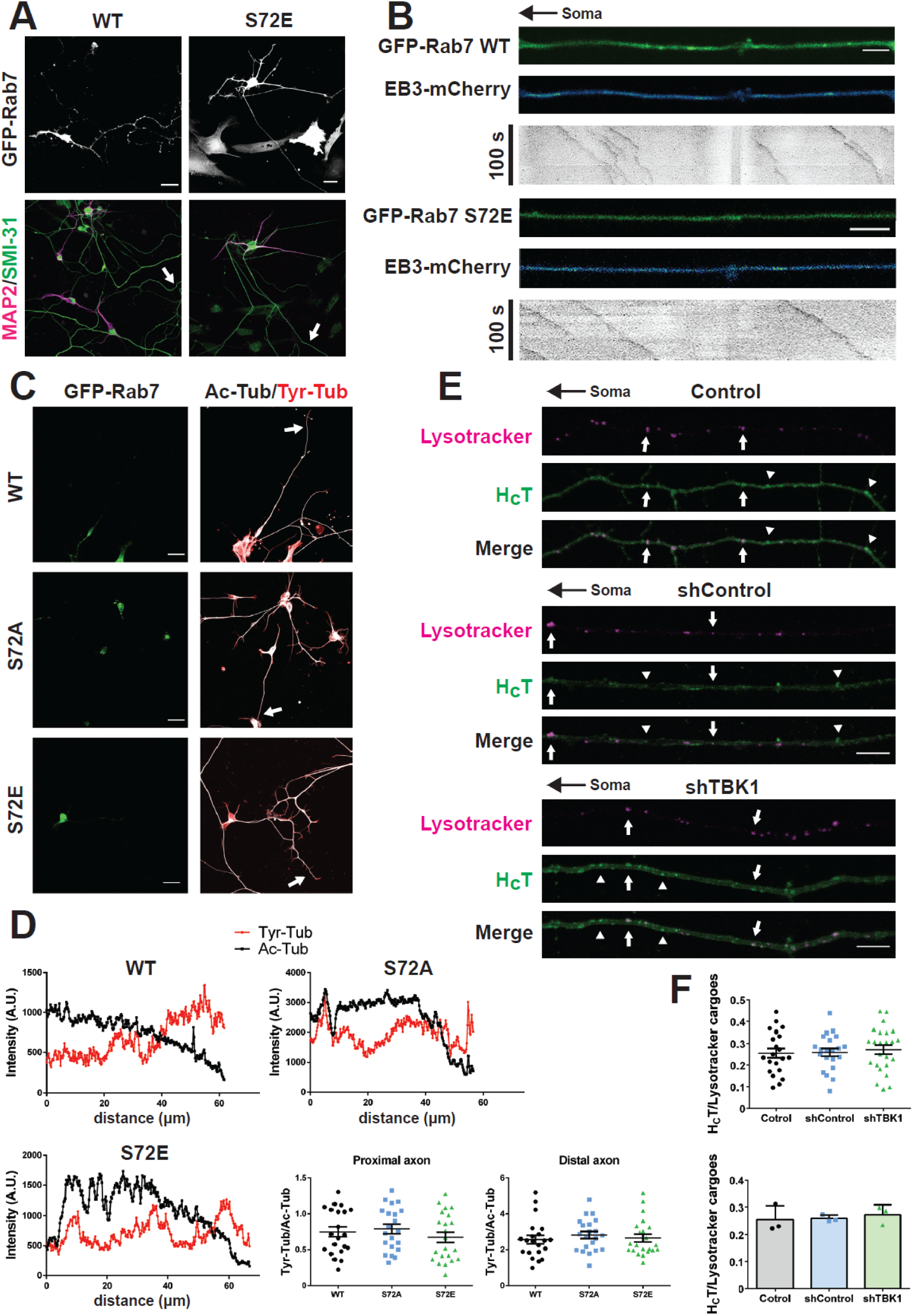
The bidirectional transport phenotype is not related to alterations in neuronal polarity or endosomal sorting defects. **(A)** Primary MNs were magnetofected on DIV 2 with either GFP-Rab7 WT or S72E, fixed after 8 h and stained as indicated. Arrows show the axon of the transfected neuron in the merged image. Magnetofected MNs acquire normal axonal and somatodendritic compartments. Scale bar, 25 μm. **(B)** Primary MNs were co-magnetofected with EB3-mCherry and either GFP-Rab7 WT or S72E on DIV 2 and imaged after 6 h. Still images from time-lapses, as well as kymographs are shown. The black arrow on top indicates the position of the soma. EB3-mCherry fluorescence signal is shown with a blue-green look-up table. Axonal microtubules are oriented with their plus-end out in both conditions. Number of retrogradely moving comets: WT first experiment = 1/39, S72E first experiment = 3/133, WT second experiment = 2/98, S72E second experiment = 1/89. Scale bar, 5 μm. **(C)** Primary MNs were magnetofected on DIV 2 with GFP-Rab7 WT, GFP-Rab7 S72A or GFP-Rab7 S72E, simultaneously fixed and extracted after 24 h, followed by immunolabelling. Arrows indicate the axon corresponding to the magnetofected neuron. Scale bar, 25 μm. **(D)** Fluorescence intensity profiles were measured starting at the axon tip until collecting profiles of at least 50 μm in length. In the fluorescence intensity profiles, the soma is located towards the left. The ratio between Tyr-tubulin/Ac-tubulin on the proximal and distal axon was determined (average measurement from segments of 5 μm in length, between 5 and 10 μm from the axon tip (distal) and between 45 and 50 μm from the axonal tip (proximal)). No change in the proximal or distal Tyr-tubulin/Ac-tubulin ratio was detected. Number of neurons: WT 21, S72A 21, S72E 22, from three independent primary cultures; one-way ANOVA with Tukey’s multiple comparison test. **(E)** Primary MNs were transduced with either shControl or shTBK1 on DIV 3. 72 h later, MNs were labelled with AlexaFluor 647-H_C_T (30 nM, 45 min) and lysotracker Red (50 nM, 30 min), washed and imaged. A set of still images from time-lapses for each condition is presented. Examples of organelles double positive for lysotracker and H_C_T are indicated by white arrows, whilst carriers labelled by H_C_T only are shown by arrowheads. The black arrow indicates the orientation of the axon. Scale bar, 10 µm. **(F)** Quantification of cargoes double positive for lysotracker and H_C_T per MN (upper panel) and per experiment (lower panel). About ∼ 25% of H_C_T-positive signalling endosomes are also labelled by Lysotracker, a percentage that remained constant for every condition. Number of neurons: Control 21, shControl 21, shTBK1 24, from three independent primary cultures); one-way ANOVA with Tukey’s multiple comparison test.

**Supplementary Figure 3.**
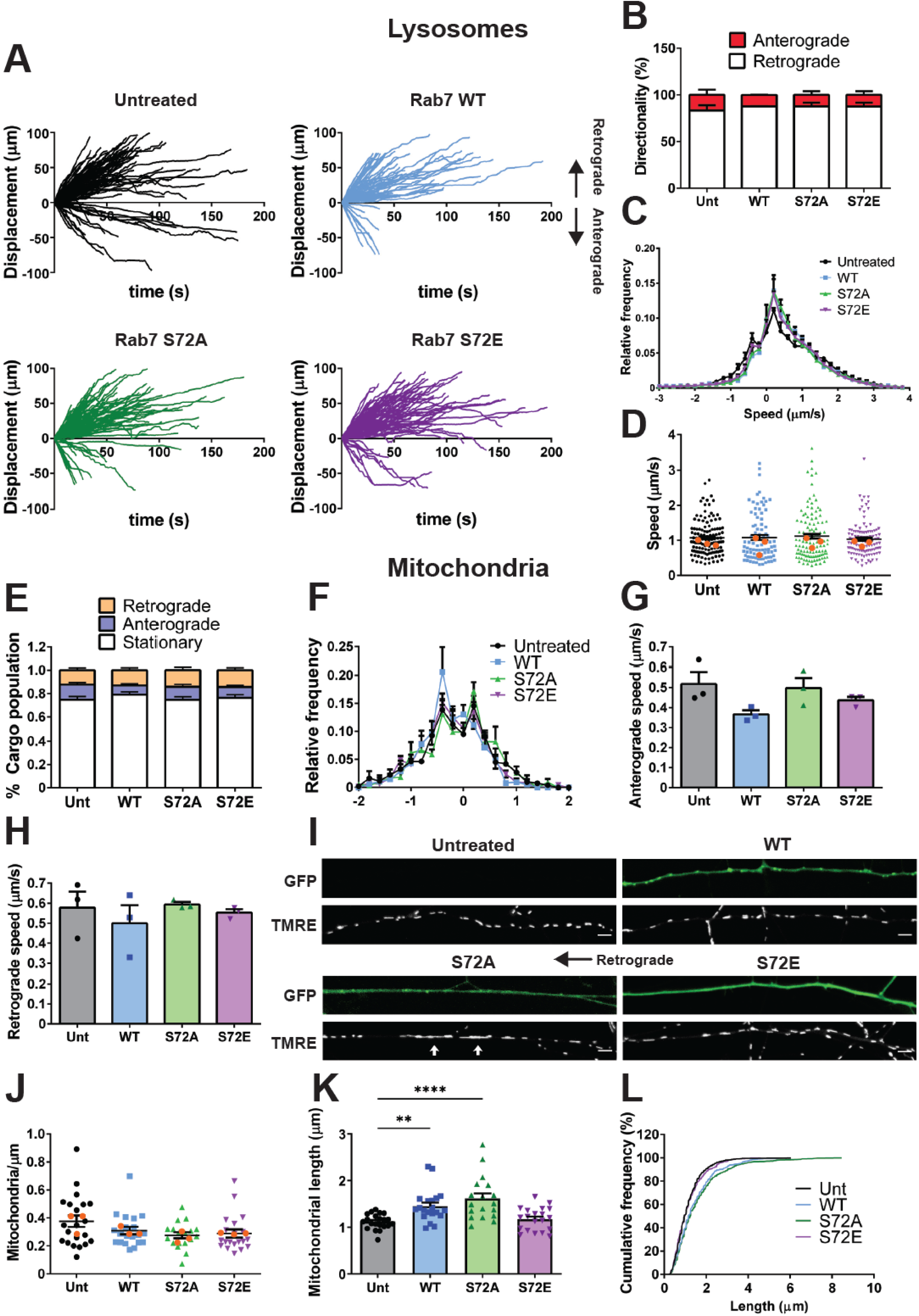
Lysosomal and mitochondrial axonal transport are not affected by Rab7 S72E expression. Primary MNs were magnetofected with either GFP-Rab7 WT, S72A or 72E on DIV 5 and 24 h later, labelled with Lysotracker Red (50 nM, 30 min), washed and imaged. **(A)** Displacement graphs for the untreated and magnetofected conditions, with retrograde transport shown as positive. Despite lysosomal bidirectional movement, carriers exhibit biased retrograde trajectories, which are not changed by Rab7 WT or the mutant variants. **(B)** Quantification of the directionality of transport. About 80% of lysotracker positive carriers move retrogradely in all conditions. n = 3, two-way ANOVA with Sidak’s multiple comparison test. **(C)** Speed profiles for every condition. Speed profiles overlap and indicate lysosomal transport is not affected by Rab S72E expression. **(D)** Average speed per cargo, considering every speed as positive. No statistically significant differences were observed. Number of cargoes: Unt 127, WT 89, S72A 117, S72E 111, from three independent primary cultures, orange dots represent the mean for each replicate. One-way ANOVA with Tukey’s multiple comparison test. **(E)** Primary MNs in culture were magnetofected with either GFP-Rab7 WT, S72A or 72E on DIV 5 and labelled 24 h later with TMRE (20 nM, 20 min). Cargo population distribution. 70-80% of mitochondria were static, whilst the rest were split equally between retrograde and anterograde directions. No significant changes were detected among conditions. Number of neurons: Unt 21, WT 20, S72A 18, S72E 20 from three independent cultures; two-way ANOVA with Sidak’s multiple comparison test. **(F)** Speed profiles for every condition. The overlapping speed profiles shows that axonal transport of mitochondria is not affected by Rab72E expression. **(G)** Average retrograde speed per experiment. Average speeds remain unchanged, although there was a reduced range for S72A and S72E, hence, a non-parametric test was used. n = 3, Kruskal-Wallis one-way ANOVA with Dunn’s multiple comparison test. **(H)** Average anterograde speed. No changes were detected between conditions. n = 3, one-way ANOVA with Tukey’s multiple comparison test. **(I)** Representative images from live-cell imaging experiments, with primary MNs expressing the indicated Rab7 constructs. White arrows show examples of elongated mitochondria in Rab7 S72A-expressing neurons. The black arrow indicates the orientation of the axon. Scale bar, 5 μm. **(J)** Mitochondrial axonal density for every condition. No statistical differences were detected between conditions. Number of neurons: Unt 21, WT 20, S72A 18, S72E 20 from three independent primary cultures; one-way ANOVA with Tukey’s multiple comparison test. **(K)** Mitochondrial length in axons. Rab7 WT induced a mild increase in mitochondrial length, which was more evident in Rab7 S72A-expressing MNs, whilst Rab7 S72E-expressing MNs remained indistinguishable from untreated controls. Number of neurons: Unt 21, WT 20, S72A 18, S72E 20 from three independent cultures; one-way ANOVA with Tukey’s multiple comparison test, **, P < 0.01, ****, P < 0.0001. **(L)** The cumulative frequency for mitochondrial length shows similar distributions for Rab7 S72E and the untreated control. Conversely, both Rab7 WT- and Rab7 S72A-expressing MNs are shifted toward higher length values.

**Supplementary Figure 4.**
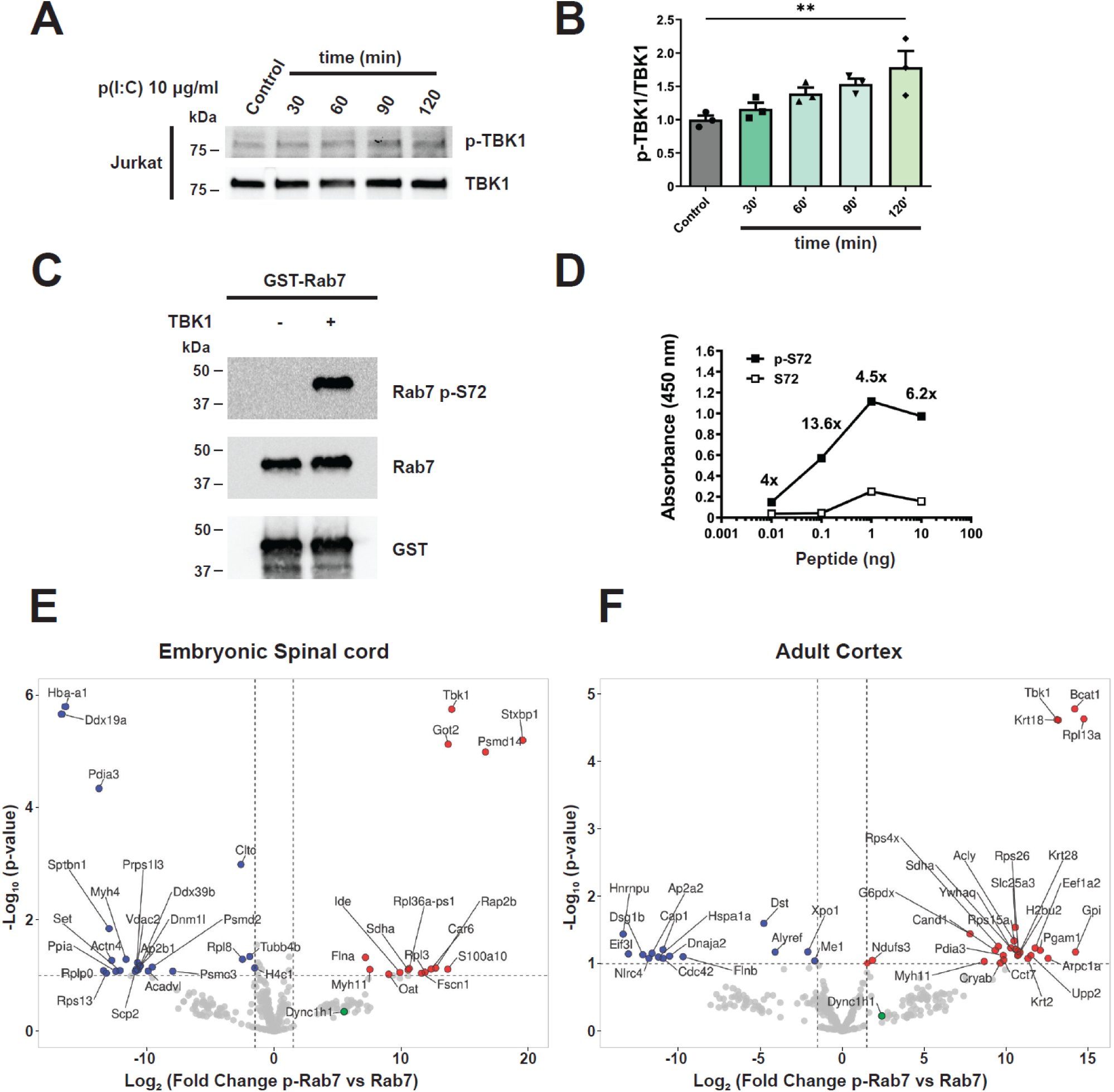
Rab7 p-S72 antibody validation and identification of differential interactors. **(A)** Jurkat cells were stimulated with 10 μg/ml p(I:C) for 30, 60, 90 or 120 min. Afte this, TBK1 p-S172 and total TBK1 levels were determined by WB. **(B)** Quantification of TBK1 p-S172/TBK1 from (A). n = 3, one-way ANOVA with Dunnett’s multiple comparison test. **P < 0.01. **(C)** WB of a TBK1 *in vitro* kinase assay using GST-Rab7 as substrate. The phosphorylation was detected with our anti-Rab7 p-S72 antibody. A Rab7 p-S72 band appears only when TBK1 is added. **(D)** ELISA to assess the binding of anti-Rab7 p-S72 to a phosphorylated or unphosphorylated peptide flanking Rab7 S72. The antibody shows specificity for the phosphorylated peptide. **(E,F)** GST-Rab7 *in vitro* phosphorylated by TBK1 (or unphosphorylated GST-Rab7 as control) were incubated with protein extracts derived from mouse embryonic spinal cord or adult cortex, respectively. The interactors were identified by mass spectrometry and are shown in the volcano plots. Red dots correspond to binding partners preferring Rab7 p-S72, blue dots represent interactors which bind preferentially to unphosphorylated Rab7. The green dot shows cytoplasmic dynein heavy chain (Dync1h1). All interactors above the threshold of -Log_10_ (p value) = 1 and Log_2_ (Fold change) = 1.5 have been labelled with their gene name.

## References

1. Sleigh JN, Rossor AM, Fellows AD, Tosolini AP, Schiavo G. Axonal transport and neurological disease. Nat Rev Neurol. 2019;15:691–703.

2. Scott-Solomon E, Kuruvilla R. Mechanisms of neurotrophin trafficking via Trk receptors. Mol Cell Neurosci. 2018;91:25–33.

3. Villarroel-Campos D, Schiavo G, Lazo OM. The many disguises of the signalling endosome. FEBS Lett. 2018;592:3615–32.

4. Deinhardt K, Salinas S, Verastegui C, Watson R, Worth D, Hanrahan S, et al. Rab5 and Rab7 control endocytic sorting along the axonal retrograde transport pathway. Neuron. 2006;52:293–305.

5. Bilsland LG, Sahai E, Kelly G, Golding M, Greensmith L, Schiavo G. Deficits in axonal transport precede ALS symptoms in vivo. Proc Natl Acad Sci USA. 2010;107:20523–8.

6. Sleigh JN, Tosolini AP, Gordon D, Devoy A, Fratta P, Fisher EMC, et al. Mice Carrying ALS Mutant TDP-43, but Not Mutant FUS, Display In Vivo Defects in Axonal Transport of Signaling Endosomes. Cell Rep. 2020;30:3655–3662.e2.

7. Tosolini AP, Sleigh JN, Surana S, Rhymes ER, Cahalan SD, Schiavo G. BDNF-dependent modulation of axonal transport is selectively impaired in ALS. Acta Neuropathol Commun. 2022;10:121.

8. Hutagalung AH, Novick PJ. Role of Rab GTPases in membrane traffic and cell physiology. Physiol Rev. 2011;91:119–49.

9. Homma Y, Hiragi S, Fukuda M. Rab family of small GTPases: an updated view on their regulation and functions. FEBS J. 2021;288:36–55.

10. Heo J-M, Ordureau A, Swarup S, Paulo JA, Shen K, Sabatini DM, et al. RAB7A phosphorylation by TBK1 promotes mitophagy via the PINK-PARKIN pathway. Sci Adv. 2018;4:eaav0443.

11. Hanafusa H, Yagi T, Ikeda H, Hisamoto N, Nishioka T, Kaibuchi K, et al. LRRK1 phosphorylation of Rab7 at S72 links trafficking of EGFR-containing endosomes to its effector RILP. J Cell Sci. 2019;132.

12. Babur Ö, Melrose AR, Cunliffe JM, Klimek J, Pang J, Sepp A-LI, et al. Phosphoproteomic quantitation and causal analysis reveal pathways in GPVI/ITAM-mediated platelet activation programs. Blood. 2020;136:2346–58.

13. Modica G, Tejeda-Valencia L, Sauvageau E, Yasa S, Maes J, Skorobogata O, et al. Phosphorylation on serine 72 modulates Rab7A palmitoylation and retromer recruitment. J Cell Sci. 2025;138:jcs262177.

14. Ritter JL, Zhu Z, Thai TC, Mahadevan NR, Mertins P, Knelson EH, et al. Phosphorylation of RAB7 by TBK1/IKKε Regulates Innate Immune Signaling in Triple-Negative Breast Cancer. Cancer Res. 2020;80:44–56.

15. Shinde SR, Maddika S. PTEN modulates EGFR late endocytic trafficking and degradation by dephosphorylating Rab7. Nat Commun. 2016;7:10689.

16. Tudorica DA, Basak B, Puerta Cordova AS, Khuu G, Rose K, Lazarou M, et al. A RAB7A phosphoswitch coordinates Rubicon Homology protein regulation of Parkin-dependent mitophagy. J Cell Biol. 2024;223:e202309015.

17. Ahmad L, Zhang S-Y, Casanova J-L, Sancho-Shimizu V. Human TBK1: A Gatekeeper of Neuroinflammation. Trends Mol Med. 2016;22:511–27.

18. Harding O, Evans CS, Ye J, Cheung J, Maniatis T, Holzbaur ELF. ALS- and FTD-associated missense mutations in TBK1 differentially disrupt mitophagy. Proc Natl Acad Sci U S A. 2021;118.

19. Cirulli ET, Lasseigne BN, Petrovski S, Sapp PC, Dion PA, Leblond CS, et al. Exome sequencing in amyotrophic lateral sclerosis identifies risk genes and pathways. Science. 2015;347:1436–41.

20. Freischmidt A, Wieland T, Richter B, Ruf W, Schaeffer V, Müller K, et al. Haploinsufficiency of TBK1 causes familial ALS and fronto-temporal dementia. Nat Neurosci. 2015;18:631–6.

21. Talaia G, Bentley-DeSousa A, Ferguson SM. Lysosomal TBK1 responds to amino acid availability to relieve Rab7-dependent mTORC1 inhibition. EMBO J. 2024;43:3948–67.

22. Shao W, Todd TW, Wu Y, Jones CY, Tong J, Jansen-West K, et al. Two FTD-ALS genes converge on the endosomal pathway to induce TDP-43 pathology and degeneration. Science. 2022;378:94–9.

23. Ye J, Cheung J, Gerbino V, Ahlsén G, Zimanyi C, Hirsh D, et al. Effects of ALS-associated TANK binding kinase 1 mutations on protein–protein interactions and kinase activity. PNAS [Internet]. 2019 [cited 2019 Dec 1]; Available from: https://www.pnas.org/content/early/2019/11/19/1915732116

24. Gurfinkel Y, Polain N, Sonar K, Nice P, Mancera RL, Rea SL. Functional and structural consequences of TBK1 missense variants in frontotemporal lobar degeneration and amyotrophic lateral sclerosis. Neurobiology of Disease. 2022;174:105859.

25. Brenner D, Sieverding K, Bruno C, Lüningschrör P, Buck E, Mungwa S, et al. Heterozygous Tbk1 loss has opposing effects in early and late stages of ALS in mice. J Exp Med. 2019;216:267–78.

26. Lalli G, Schiavo G. Analysis of retrograde transport in motor neurons reveals common endocytic carriers for tetanus toxin and neurotrophin receptor p75NTR. J Cell Biol. 2002;156:233–9.

27. Maday S, Wallace KE, Holzbaur ELF. Autophagosomes initiate distally and mature during transport toward the cell soma in primary neurons. J Cell Biol. 2012;196:407–17.

28. Cason SE, Carman PJ, Van Duyne C, Goldsmith J, Dominguez R, Holzbaur ELF. Sequential dynein effectors regulate axonal autophagosome motility in a maturation-dependent pathway. J Cell Biol. 2021;220.

29. Ferguson SM. Neuronal lysosomes. Neurosci Lett. 2019;697:1–9.

30. Mandal A, Drerup CM. Axonal Transport and Mitochondrial Function in Neurons. Front Cell Neurosci. 2019;13:373.

31. Czerkies M, Korwek Z, Prus W, Kochańczyk M, Jaruszewicz-Błońska J, Tudelska K, et al. Cell fate in antiviral response arises in the crosstalk of IRF, NF-κB and JAK/STAT pathways. Nat Commun. 2018;9:493.

32. Pillai S, Nguyen J, Johnson J, Haura E, Coppola D, Chellappan S. Tank binding kinase 1 is a centrosome-associated kinase necessary for microtubule dynamics and mitosis. Nat Commun. 2015;6:10072.

33. Johnson DE, Ostrowski P, Jaumouillé V, Grinstein S. The position of lysosomes within the cell determines their luminal pH. J Cell Biol. 2016;212:677–92.

34. Cantalupo G, Alifano P, Roberti V, Bruni CB, Bucci C. Rab-interacting lysosomal protein (RILP): the Rab7 effector required for transport to lysosomes. EMBO J. 2001;20:683–93.

35. Johansson M, Rocha N, Zwart W, Jordens I, Janssen L, Kuijl C, et al. Activation of endosomal dynein motors by stepwise assembly of Rab7-RILP-p150Glued, ORP1L, and the receptor betalll spectrin. J Cell Biol. 2007;176:459–71.

36. Zhou B, Cai Q, Xie Y, Sheng Z-H. Snapin recruits dynein to BDNF-TrkB signaling endosomes for retrograde axonal transport and is essential for dendrite growth of cortical neurons. Cell Rep. 2012;2:42–51.

37. Olenick MA, Dominguez R, Holzbaur ELF. Dynein activator Hook1 is required for trafficking of BDNF-signaling endosomes in neurons. J Cell Biol. 2019;218:220–33.

38. Wong YC, Ysselstein D, Krainc D. Mitochondria-lysosome contacts regulate mitochondrial fission via RAB7 GTP hydrolysis. Nature. 2018;554:382–6.

39. Dou D, Smith EM, Evans CS, Boecker CA, Holzbaur ELF. Regulatory imbalance between LRRK2 kinase, PPM1H phosphatase, and ARF6 GTPase disrupts the axonal transport of autophagosomes. Cell Rep. 2023;42:112448.

40. Brenner D, Sieverding K, Srinidhi J, Zellner S, Secker C, Yilmaz R, et al. A TBK1 variant causes autophagolysosomal and motoneuron pathology without neuroinflammation in mice. J Exp Med. 2024;221:e20221190.

41. Malik AU, Karapetsas A, Nirujogi RS, Mathea S, Chatterjee D, Pal P, et al. Deciphering the LRRK code: LRRK1 and LRRK2 phosphorylate distinct Rab proteins and are regulated by diverse mechanisms. Biochem J. 2021;478:553–78.

42. Debaisieux S, Encheva V, Chakravarty P, Snijders AP, Schiavo G. Analysis of Signaling Endosome Composition and Dynamics Using SILAC in Embryonic Stem Cell-Derived Neurons. Mol Cell Proteomics. 2016;15:542–57.

43. Guardia CM, Farías GG, Jia R, Pu J, Bonifacino JS. BORC Functions Upstream of Kinesins 1 and 3 to Coordinate Regional Movement of Lysosomes along Different Microtubule Tracks. Cell Rep. 2016;17:1950–61.

44. Boecker CA, Goldsmith J, Dou D, Cajka GG, Holzbaur ELF. Increased LRRK2 kinase activity alters neuronal autophagy by disrupting the axonal transport of autophagosomes. Curr Biol. 2021;31:2140–2154.e6.

45. Cason SE, Holzbaur ELF. Axonal transport of autophagosomes is regulated by dynein activators JIP3/JIP4 and ARF/RAB GTPases. J Cell Biol. 2023;222:e202301084.

46. Vides EG, Adhikari A, Chiang CY, Lis P, Purlyte E, Limouse C, et al. A feed-forward pathway drives LRRK2 kinase membrane recruitment and activation. Elife. 2022;11:e79771.

47. Goethals S, Ydens E, Timmerman V, Janssens S. Toll-like receptor expression in the peripheral nerve. Glia. 2010;58:1701–9.

48. Ye J, Dephoure N, Maniatis T. Complex Intracellular Mechanisms of TBK1 Kinase Activation Revealed by a Specific Small Molecule Inhibitor [Internet]. 2022 [cited 2025 Apr 10]. Available from: http://biorxiv.org/lookup/doi/10.1101/2022.10.11.511671

49. Hao J, Wells MF, Niu G, San Juan IG, Limone F, Fukuda A, et al. Loss of TBK1 activity leads to TDP-43 proteinopathy through lysosomal dysfunction in human motor neurons [Internet]. 2021 [cited 2025 Apr 10]. Available from: http://biorxiv.org/lookup/doi/10.1101/2021.10.11.464011

50. Bucci C, Thomsen P, Nicoziani P, McCarthy J, van Deurs B. Rab7: a key to lysosome biogenesis. Mol Biol Cell. 2000;11:467–80.

51. Arce V, Garces A, de Bovis B, Filippi P, Henderson C, Pettmann B, et al. Cardiotrophin-1 requires LIFRbeta to promote survival of mouse motoneurons purified by a novel technique. J Neurosci Res. 1999;55:119–26.

52. Goedhart J, Luijsterburg MS. VolcaNoseR is a web app for creating, exploring, labeling and sharing volcano plots. Sci Rep. 2020;10:20560.

53. Ahmad FJ, Hughey J, Wittmann T, Hyman A, Greaser M, Baas PW. Motor proteins regulate force interactions between microtubules and microfilaments in the axon. Nat Cell Biol. 2000;2:276–80.

54. Schindelin J, Arganda-Carreras I, Frise E, Kaynig V, Longair M, Pietzsch T, et al. Fiji: an open-source platform for biological-image analysis. Nat Methods. 2012;9:676–82.

55. Tinevez J-Y, Perry N, Schindelin J, Hoopes GM, Reynolds GD, Laplantine E, et al. TrackMate: An open and extensible platform for single-particle tracking. Methods. 2017;115:80–90.

56. Neumann S, Chassefeyre R, Campbell GE, Encalada SE. KymoAnalyzer: a software tool for the quantitative analysis of intracellular transport in neurons. Traffic. 2017;18:71–88.

